# Human chronic inflammation is orchestrated by spatially restricted inflammation-activated Dendritic cells

**DOI:** 10.1101/2025.05.19.654443

**Authors:** J.R. Thomas, J. Haub, A. Hunecke, J. Yu, P. Maier, M. Zhang, B.E.F. Pregler, M. Raspe, S. Strieth, M. Schneider, V. Schäfer, C. Pizzaro, D. Skowasch, F. Ginhoux, M. Toma, J. Martin, T. Send, A. Schlitzer

## Abstract

Dendritic cells (DCs) form coordinated networks that orchestrate inflammatory responses across tissues. Upon activation, conventional dendritic cells (DCs) undergo changes in their transcriptome, phenotype, and function, depending on the tissue microenvironment. The role of DC activation during human chronic inflammation has yet to be explored. To address this, we aimed to investigate activated DCs across a range of chronically inflamed human tissues, with a focus on cervical lymph nodes (LNs). We performed scRNAseq and flow cytometry on healthy, acutely, chronically inflamed, and cancer-associated human cervical (LNs) and identified a novel inflammation-associated DC (iaDC) cell state, found only within chronically inflamed LNs. iaDCs were defined by a unique cytokine expression profile (*CXCL9, CXCL10, IL1B*) and could be phenotypically distinguished from other activated DCs by their elevated expression of CD1C, CD206, CD319, and CD274. Functionally, we found that IFNγ signaling induces DC2s to enter the iaDC cell state and that this process can be abrogated by inhibition of JAK-STAT signaling. Using spatial transcriptomics, we observed LN iaDC residing within a specific chronic inflammatory niche, which was enriched for inflammatory monocytes, NK cells, and effector memory CD4^+^ and CD8^+^ T cells. Extrapolating our findings to other chronic inflammation-associated diseases, we similarly observed the emergence of iaDCs within the intestines of treatment-resistant Crohn’s Disease patients, synovial membranes and lymph nodes of rheumatoid arthritis patients and lungs of sarcoidosis patients. Similarly to the lymph node, iaDCs were found to reside in conserved chronic inflammation-associated spatial niches within the small intestine of affected individuals and displayed equivalent transcriptional characteristics and surrounding cellular neighborhoods. Collectively, these data highlight a previously unexplored role of iaDCs in human chronic inflammatory diseases and propose a conserved spatially restricted iaDC-populated chronic inflammatory niche associated with resistance to therapy. These findings highlight novel avenues to shape inflammatory trajectories by targeting iaDCs and their spatial niches, for example, by inhibiting JAK-STAT signaling.

## Introduction

Immune-mediated inflammatory diseases (IMIDs) comprise a heterogenous group of disorders affecting various organs, characterised by sustained inflammation resulting in tissue damage and organ dysfunction ^1–3^. Although substantial progress has been made in our understanding the cellular and molecular determinants of IMIDs in recent years, the pathobiology of (non)response to therapy is not well understood, and a significant proportion of patients (30-40% in Crohn’s Disease and Rheumatoid Arthritis) fail to respond to gold-standard therapeutic intervention ^4–6^.

Dendritic cells (DCs) have critical roles in the initiation, propagation and resolution of immune responses, due to their capacity to sense their microenvironment, migrate to lymphoid structures and orchestrate T cell responses ^7^. Recent work has revealed that both conventional DC1 and DC2 subsets can be induced to enter an activated DC (aDC) cell state, characterized by a highly conserved phenotypic, transcriptional, and functional program ^8–12^. This cell state has been variably named as “activated DC”, “mature DC”, “migratory DC”, “CCR7+ DC”, “LAMP3+ DC” and “mature DCs enriched in immunoregulatory molecules (mregDC)”. Recent work has begun to decipher context-specific aDC programmes, for example homeostatic DC activation governed by the uptake of apoptotic cells ^8,12^. However, our understanding of how context and niche-specific signalling cues induce the acquisition of diverse DC activation states, and how these states initiate and perpetuate inflammation, remains limited.

IMIDs are often characterised by the accumulation of diverse pathogenic effector memory T cell populations, which exhibit clonal expansion, cytotoxicity and pro-inflammatory cytokine secretion ^13–19^. The instruction of T cells towards these pathogenic states in IMIDS is, in turn, dependent on the functional properties of interacting aDCs. However, it remains unclear which specific intratissular cues drive the development of DC activation states capable of inducing these pathogenic T cell populations and thus chronic inflammation. To better understand the determinants of disease trajectories and their connection towards therapy-responsiveness in IMIDs we require a deeper insight into the functional and spatial regulation of DC activation in chronic inflammatory settings.

In this work we leveraged scRNAseq, spatial transcriptomics and *in vitro* functional analyses to investigate the emergent properties of aDCs during chronic inflammation in humans. We identified a population of inflammatory aDC (iaDCs) which arise during chronic inflammation of human lymph nodes in an IFN-dependent fashion, and occupy a specific chronic inflammatory niche. Crucially, iaDCs induce the further expression of IFNγ in CD8+ T cells *in vitro*, suggesting that iaDCs orchestrate chronic inflammation by guiding pathogenic T-cell phenotypes. Finally, we demonstrate that iaDCs, along with their associated niche, are a conserved feature of various IMIDs. Collectively, these findings highlight a conserved role of iaDCs in the pathogenesis of IMIDs, and draw attention to future possibilities of shaping inflammatory trajectories via therapeutic targeting of iaDCs and their niches.

## Results

### Chronic inflammation drives a specific dendritic cell activation state in human lymph nodes

Dendritic cell (DC) activation in humans is a plastic process, dependent on an array of signalling cues available within the spatial microenvironments DCs reside. DC activation induced by acute sterile or infectious agent-elicited inflammation, and during cancer, has been extensively studied. However, the molecular determinants of DC activation states during human chronic inflammation remain largely elusive. Thus, to better understand the emergent properties of DC activation states during human chronic inflammation, we first characterized the abundance of human CCR7^+^ DCs in chronically inflamed cervical lymph nodes (LNs), bronchoalveolar and synovial fluids, subcutaneous adipose tissues, and peripheral blood mononuclear cells (PBMC) using flow cytometry (Figure 1A, B S1). This analysis showed that activated CCR7^+^ DCs were present in all analysed tissues, with the exception of the PBMC, in individuals affected by chronic human inflammatory conditions (Figure 1A, B). Acute and chronic swelling of cervical LNs is common, and if severe, is resolved by surgical resection of the affected LN. Thus, acute and chronically inflamed cervical LNs provide a unique opportunity to dissect the molecular and cellular underpinnings of human chronic inflammation. To study these processes, we collected surgically resected cervical LNs which were acutely (≤ 4 weeks) or chronically inflamed ( > 3 month) or lymphoma-associated. Inflamed LN donors were otherwise healthy and negative for all testable infectious disease associated parameters as well as cancer.

**Figure 1.**
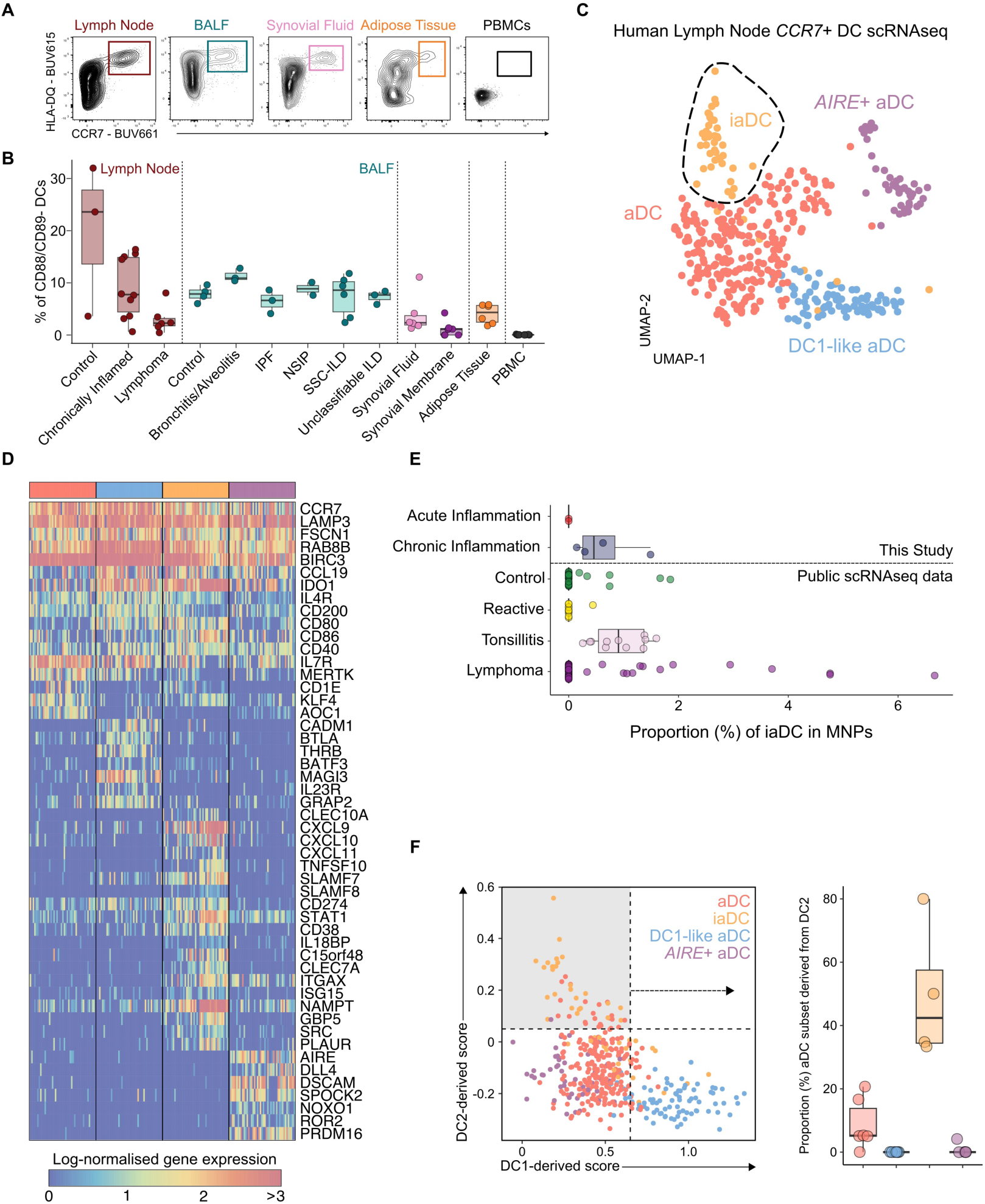
iaDC emerge during chronic inflammation of human lymph nodes. **A)** Flow cytometry plots of HLA-DQ and CCR7 expression in DCs across indicated human tissues. **B**) Quantification of CCR7+ DC abundance within CD88/CD89-DCs across indicated human tissues. **C**) UMAP visualisation of scRNAseq of CCR7+ DC from human LNs. aDC-Activated DC, iaDC – inflammatory activated DC. *n* = 7. **D**). Heatmap depicting gene expression of conserved and differentially expressed genes in CCR7+ DC subclusters. **E**). Quantification of iaDC abundance in integrated scRNAseq data, stratified by sample type. **F**) Biaxial plot of enrichment scores for DC1- and DC2-derived gene signatures from (Cheng 2021) in CCR7+ DC scRNAseq data, and (right) quantification of the proportion of each subcluster falling within the DC2-derived “gate” as indicated by the shaded area on the enrichment plot.

To characterize the transcriptional underpinnings of human chronic inflammation associated DC activation we depleted B and T-cells from our LN preparations and generated single cell transcriptomic data of LN-resident mononuclear phagocytes (Figure S2A-D). Next, we focused our analysis on CCR7^+^ DCs (Figure 1C), revealing 4 transcriptionally distinct clusters of activated CCR7^+^ DCs within human cervical LNs. All clusters expressed prototypic DC activation markers including high expression of *CCR7*, *LAMP3*, *FSCN1* and *BIRC3*. We denoted a cluster expressing *CD1E*, *KLF4* and AOC1 as activated DC (aDC) (red), and a cluster enriched for DC1-related transcripts *CADM1*, *BATF3* and *BTLA* as DC1-like aDC (Blue). A third cluster highly expressed inflammatory gene s including *CXCL9*, *CXCL10, CXCL11*, *ISG15*, *STAT1*, *SLAMF7* and *CD274* (PD-L1) and as such, was termed inflammatory aDC (iaDC) (yellow) (Figure 1D). Lastly, a cluster with high expression of *AIRE*, *DLL4*, *DSCAM* and *PRDM16*, transcriptionally similar to Janus Cells described by others, (Figure 1D) was labelled here as *AIRE+* aDC (violet). Next, we visualized the abundance of CCR7^+^ DC transcriptional subsets across our disease conditions (Figure S2E,F). These data showed that iaDCs were overrepresented in chronically inflamed LNs, and iaDC abundance did not strongly correlate with other CCR7^+^ DC clusters, suggesting a specific emergence of iaDCs in this condition. To validate the iaDC subset at the protein-level, we identified genes upregulated in iaDCs encoding surface markers (Fig S2G) and analysed their expression in CCR7+ DCs in chronically inflamed LNs by flow cytometry. We identified a phenotypic subset of CCR7+ DCs consistent with iaDCs expressing CD319(*SLAMF7*), CD274, CD86, CD11c, TRAIL and CD1c.

To further validate our scRNAseq findings, we extended our analyses by integrating our newly generated data with publicly available scRNAseq data from healthy controls, reactive (acutely inflamed) LNs, chronically inflamed tonsils (tonsilitis), and lymphoma-associated LNs (Figure 1E, S3). Here, similarly, an enrichment of chronic iaDC was detected only in tonsilitis samples but not in control, reactive, or lymphoma samples.

Next, we investigated the developmental origin of the four newly identified CCR7^+^DC subsets (Figure 1F). Here, DC1-like aDC showed a strong enrichment towards a transcriptomic DC1-derived signature, whereas aDC and *AIRE+* aDC clusters showed no enrichment towards either DC1 or DC2 origins. Lastly, iaDC showed enrichment towards a transcriptomic DC2-derived signature, suggesting a DC2 origin of this cell state prior to activation.

### IFNγ stimulates chronic inflammation-induced dendritic cell activation

To understand which molecular signals, underlie the initiation of the specific chronic inflammation-associated cellular state of iaDC we utilized NicheNet receptor ligand inference (Figure 2A). NicheNet analysis of iaDC scRNA-seq data predicted *IFNG* (encoding IFNγ) and *IFNB1* (encoding IFNβ) as regulators of the transcriptional identity of iaDC. At the pathway level further analysis confirmed ‘Response to Type II interferon’ and ‘JAK-STAT’ pathways to be specifically active in iaDC (Figure 2B, C, S4A, B). The JAK-STAT pathway is a critical constituent of IFN−driven cellular activation. To test the *ex vivo* relevance of the JAK-STAT pathway and the involvement of IFNγ in priming the iaDC cell state we utilized the clinically approved JAK-STAT inhibitor Upadacitnib (UPA) (Figure 2D). To do so we cultured cell suspensions from cervical LNs preparations, and treated cells with either 50ng/ml IFNγ, 1μm UPA, or a combination of the two, for 19h and subsequently analysed the mononuclear phagocyte compartment by flow cytometry (Figure 2E-G, S4C-F). We could identify iaDCs in *ex vivo* cultures via their CCR7^+^CD319^+^CD274^+^CXCL9^+^CXCL10^+^ phenotype (Figure 2E, F). Comparisons across culture conditions revealed that UPA treatment reduced endogenously-present iaDC upon treatment and that IFNγ, exposure significantly expands the iaDC compartment within *ex vivo* cultured cervival LNs (Figure 2G). Further co-treatment of IFNγ and UPA abrogated the IFNγ-induced expansion of iaDCs *ex vivo* (Figure 2G). To understand whether or not the environment of the cervical LN is instructive for iaDC transcriptional priming we FACS-purified PBMC-derived DC2 and treated them for 18h with IFNγ, UPA or a combination thereof (Figure 2H, S4G). Key functional iaDC-expressed molecules, such as CXCL9 and CXCL10 were specifically co-expressed upon stimulation with IFNγ, and abrogated by UPA co-treatment (Figure 2I, J, S4H) and a similar induction of iaDC-specific surface markers, such as CD274, CD319 and CD273 was observed (Figure 2K, S4H). Taken together, these data show that IFNγ imprints a JAK-STAT pathway-dependent transcriptional program on human activated DC2, specifically observed in chronically inflamed cervical LNs.

**Figure 2.**
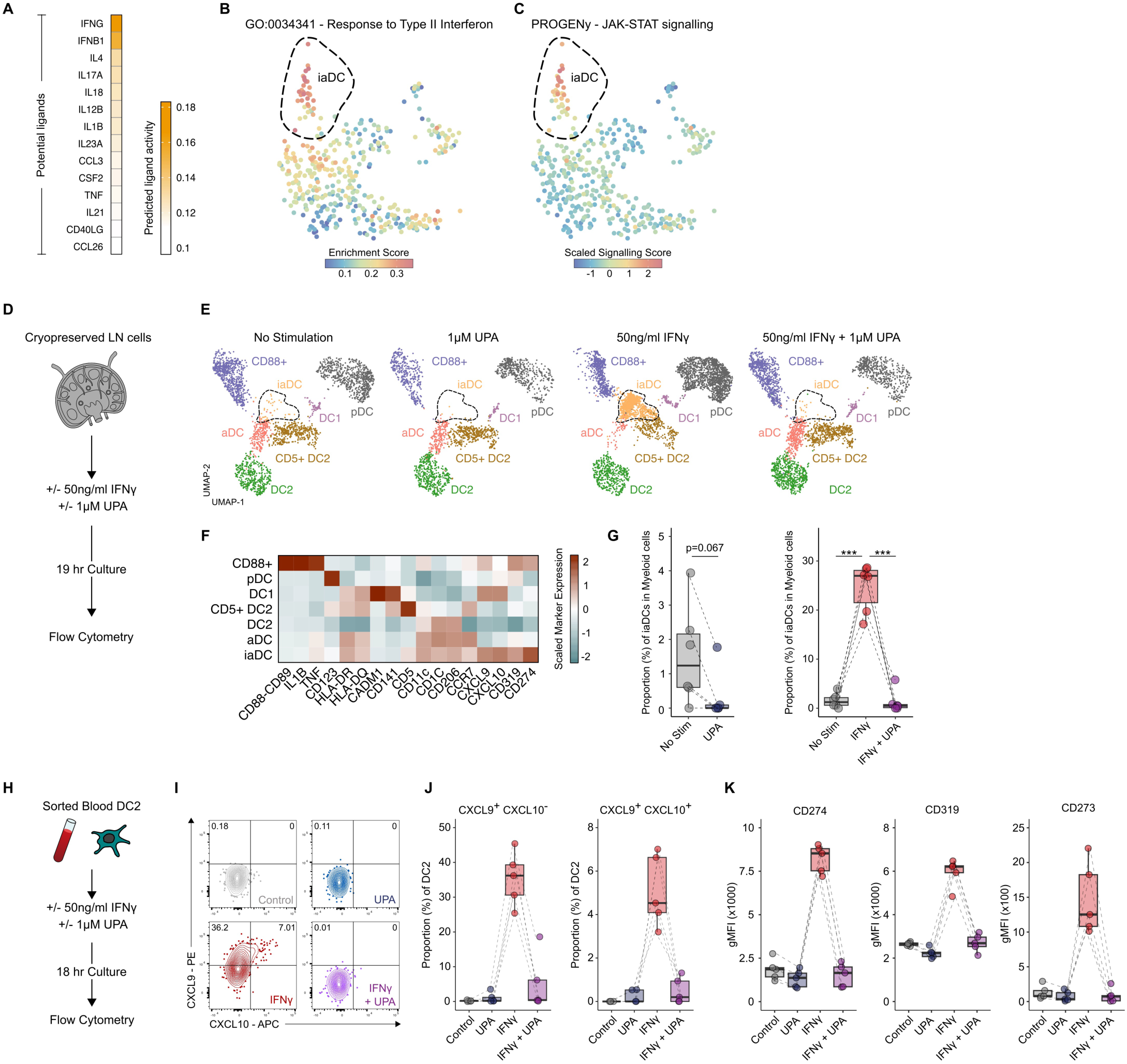
iaDC are induced by IFNγ and dependant on JAK-STAT signalling. **A)** Heatmap of predicted upstream regulators of the iaDC cell state, as determined by NicheNet analysis. Heatmap overlay of **B**) for Response to Type II Interferon gene signature enrichment score and **C**) JAK-STAT signalling score, as determined by PROGENy analysis. **D**) Schematic representation of experimental design of LN bulk stimulation experiments. UPA – Upadacitinib. **E)** UMAP visualisation of flow cytometry data from LN Mononuclear Phagocytes after stimulation. iaDCs cluster is highlighted by dotted line. **F**) Heatmap depicting phenotypes of observed clusters by flow cytometry. **G**) Quantification of iaDC abundance across experimental conditions. *n* = 6 donors. ***p > 0.001. **H**) Schematic representation of experimental design of PBMC DC2 stimulation experiments. **I**) Flow cytometry plots of CXCL9 and CXCL10 expression in DC2 after stimulation and quantification of L) CXCL9+ and CXCL19+CXCL10+ cells and **J**) CD274 (PDL1), CD319 (SLAMF7) and CD273 (PDL2) expression *n* = 5 donors.

### iaDC exist in spatially restricted niches within human cervical lymph nodes

LNs are spatially compartmentalised and architecturally heterogenous, comprising distinct anatomical regions, including the capsule, medulla, paracortex and cortex. Typically, DCs localise at the border of and within the T-cell zones to facilitate efficient T-cell activation. To investigate if activated DC states occupy distinct niches in health, acute inflammation and chronic inflammation we analysed the spatial topology of human cervical LNs using *in situ* padlock probe-based spatial transcriptomics (Figure 3A) (Xenium, 10x Genomics). We used a 477-plex probe panel, containing 100 custom probes informed by our scRNAseq analysis (Figure S5A, B), which allowed for the identification of 76 clusters across the Myeloid, T/NK cell, B cell, Stromal and Endothelial cell lineages within 8 samples (Figure S5C-G, S6).

**Figure 3.**
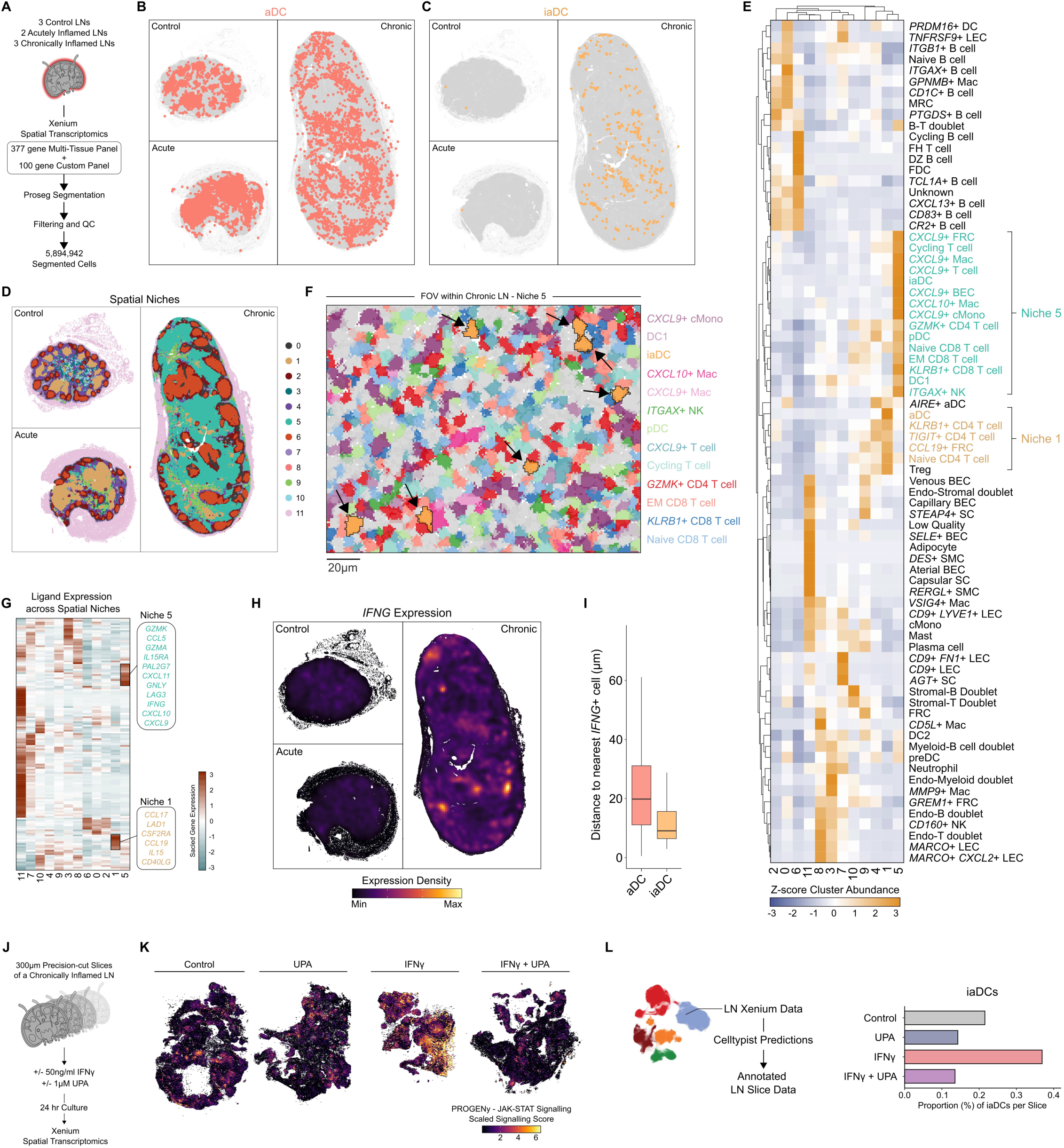
iaDC occupy a distinct spatial niche in chronically inflamed LNs. **A)** Schematic representation of Xenium experimental and pre-processing workflow. Visualisation of **B**) aDCs and **C**) iaDCs across 3 representative LN Xenium samples. **D**) Visualisation of spatial niches, as determined by 50um raster scan from each cell, across 3 representative LN Xenium samples. **E**) Heatmap depicting enrichment of Xenium clusters across calculated spatial niches. Clusters enriched in Niche 5 (iaDC-containing) and Niche 1 (aDC-containing) are highlighted. **F**) Spatial plot depicting Proseg segmented cells within a field of view (FOV) within Niche 5. Cells within clusters enriched in Niche 5 are highlighted and iaDCs are indicated with black arrows. **G**) Heatmap depicting scaled gene expression of secreted ligands across spatial niches and **H**) Visualisation of *IFNG* expression across 3 representative LN Xenium samples. **I**) Quantification of the distance from each aDC and iaDC to the nearest *IFNG*+ cell across the Xenium dataset. **J**) Schematic representation of experimental design of LN slice culture Xenium experiments. **K**) Visualisation of JAK-STAT signalling scores across LN slices, as determined by PROGENy analysis. **L**) Quantification of Celltypist-predicted iaDC abundance across LN slices.

Analysis of all activated DCs (aDCs) across control, acute, and chronically LNs showed an expected distribution of aDCs within T-cell zones (Figure 3B). However, examining the topographical distribution of iaDC within our assessed LNs revealed that within healthy and acutely inflamed LNs, very few iaDCs could be detected, in accordance with our previous scRNAseq data. However, within chronically inflamed LNs, iaDC formed clusters and their topological distribution markedly differed from the distribution observed for all aDC (Figure 3C). To further understand the topological enrichment of iaDCs within the chronically inflamed LN, we performed a spatial niche analysis of healthy, acutely, and chronically inflamed LNs (cell centroid, 50µm, Figure 3D, E S7A, B), and examined the abundance of niches across inflammatory conditions (Figure S7C) and the enrichment of cell clusters across these niches (Figure 3E, S7D). This revealed that within control and acutely inflamed LNs, typically LN-associated structures were detected, including B-cell follicles (Niche 6) close to the capsule within the cortical region, and T-cell zones (Niche 1), to which aDCs localized. Within Niche 1, typical constituents of T-cell activation were identified, including DC, Naïve CD4^+^ T-cells, and CCL19^+^ FRCs (Figure 3E). However, a different picture emerged within chronically inflamed cervical LNs.

Although B-cell-related cellular architecture was largely conserved, almost no canonical T-cell zones were detected. Instead, a new niche associated with chronic inflammation emerged, designated as Niche 5. Niche 5 was exclusively detected in chronically inflamed cervical LNs (Figure S7C) and was characterized by the presence of iaDC alongside *CXCL9*^+^ monocytes and macrophages, activated NK cells, and effector memory (EM) CD4^+^ and CD8^+^ T-cells (Figure 3E). To better understand the relevance of iaDC for the cellular architecture of Niche 5, we assessed cellular neighbours of iaDC in Niche 5 and found that iaDC are closely associated with either CD4^+^ (*GZMK*+ CD4 T cell) and CD8^+^ (EM CD8 T cell) EM T-cells (Figure 3F). Ex vivo cultures of chronically inflamed LNs indicated IFNγ as a crucial player for the transcriptional imprinting of iaDCs (Figure 2). To translate those findings to the spatial level and to reveal the topographical dependencies of the induction of the iaDC transcriptional program *in vivo* we first profiled the expression of secreted ligands within detected spatial niches across the complete spatial transcriptomic dataset (Figure 3G). Within Niche 1, we detected expression of molecules described prior to be associated to T-cell activation, such as *CCL19*, *CCL17*, *CD40LG* and *IL15*, whereas Niche 5 was enriched for molecules previously associated to chronic T-cell activation within peripheral organs, such as *GZMK*, *GZMA* and *CCL5*, alongside chemokines, such as CXCL9, CXCL10 and CXCL11, important for recruitment of EM T-cells. In line with these findings and the interferon-inducible characteristic of CXCL9, CXCL10 and CXCL11 we also found strong expression of *<u>IFNG</u>* and activation of JAK-STAT signalling within Niche 5 (Figure 3H, S7E). Further comparing the topographical association of aDC and iaDC towards *IFNG*+ cells within the chronically inflamed LN, iaDC associated closely to *IFNG* producers, such as CD4^+^ (*GZMK*+ CD4 T cell), CD8^+^ (EM CD8 T cell, *KLRB1*+ CD8 T cells) EM T-cells and *ITGAX*^+^ NK cells (Figure 3I, S7F).

To directly observe the effects of IFNγ signalling and activation of the JAK-STAT pathway in topologically intact LN sections, we utilized precision-cut slices of chronically inflamed cervical LNs, cultured with IFN-γ, UPA, or both for 24 hours, followed by spatial transcriptomic analysis (Figure 3J, S8, 10x Genomics, Xenium). Here, spatial analysis charting the activation of the JAK-STAT pathway revealed that UPA treatment significantly inhibited endogenous activation of the JAK-STAT pathway and thus resulted in reduced iaDC numbers, whereas IFNγ treatment resulted in JAK-STAT pathway activation and an increase in iaDCs, an effect was abrogated by simultaneous treatment with UPA (Figure 3K, L, S8E-K). Hence, similar to *ex vivo* LN or blood-derived DC cultures (Figure 2D-G), precision-cut LN slide cultures confirm IFNγ as a crucial molecular determinant of iaDC transcriptional programming. Together, these spatial transcriptional analysis shows that chronic inflammation rearranges the cellular configuration of LN resident niches, creating iaDC-enriched niches prone to harbor EM T-cell phenotypes.

### iaDCs drive effector T-cell responses within their niches via CXCL9-CXCR3 interaction

Spatial transcriptional analysis revealed that iaDC-dominated niches during chronic inflammation (Niche 5) are enriched for EM CD4 and CD8 T-cell phenotypes alongside NK cells (Figure 3E, S9A). To better understand this interaction, we compared the local composition of the T-cell / NK cell compartment present in Niche 1 and Niche 5 (Figure 4A, B). This revealed an increase of EM T-cells (*GZMK*^+^ CD4^+^ T-cells, *GZMK*+ CD8^+^ T-cells, *KLRB1*^+^ CD8 T-cells) and *CXCL9*^+^ T-cells within Niche 5, whereas naïve CD4^+^ T-cells and regulatory T-cells were enriched within Niche 1 (Figure 4A, B). Our prior data showed that iaDCs express high levels of *CXCL9*, and the CXCL9:CXCR3 axis is implicated in the attraction and maintenance of CD8^+^ EM T-cells (Figure 1D). To better understand the spatial organisation of EM T-cells and iaDC neighborhoods, we first mapped expression of CXCR3 within both niche 1 and 5 resident T-cell constituents (Figure 4C). Here only T-cell phenotypes enriched in Niche 5 expressed CXCR3, complementing the elevated expression of CXCL9 observed in Niche 5 (Figure 3G).

**Figure 4.**
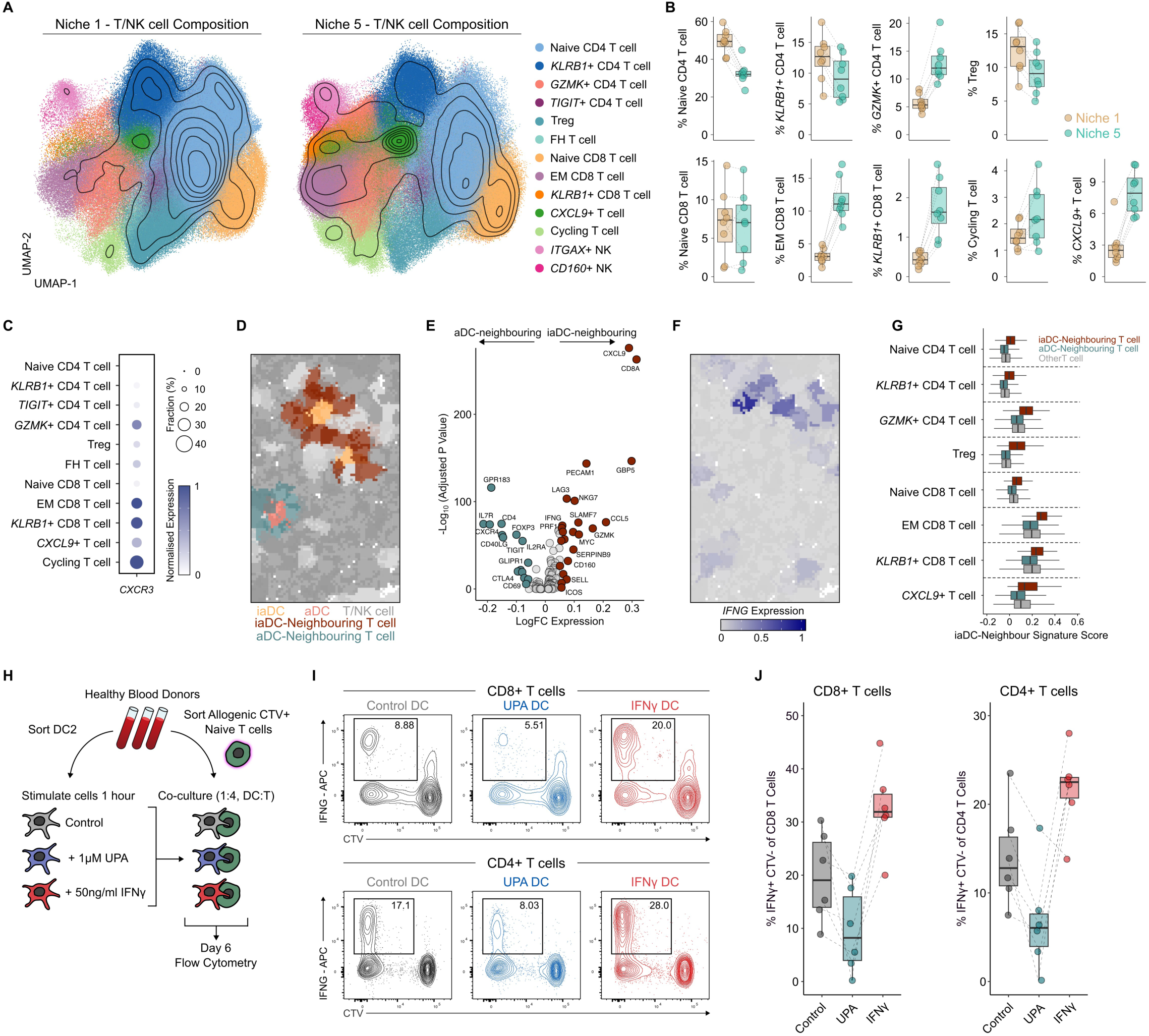
iaDC promote chronic inflammation via an IFNγ-dependant feed-forward loop. **A)** UMAP visualisation with overlain density contours of Xenium T/NK cells in Niche 1 and Niche 5 and **B**) quantification of cluster abundances per niche. **C**) Dotplot depicting normalised expression of *CXCR3* across Xenium T cell clusters. **D**) Spatial plot depicting Proseg segmented cells, highlighting aDC, iaDC and neighbouring T cell populations. **E**) Volcano plot of DEGs (logFC > 0.05 and < −0.05, adjusted p value <0.05, Wilcoxon rank sum test) between all aDC-neighbouring and iaDC-neighbouring T cells across the Xenium dataset. **F**) Boxplot depicting enrichment scores of the iaDC-neighbouring gene signature defined in E, across all T cell clusters, split by DC-neighbour status. **G**) Spatial plot as in D, but with *IFNG* expression overlain onto segmented cells. **H**) Schematic representation of experimental design of Naïve T cell stimulation experiments. **I**) Flow cytometry plots depicting IFNγ expression and proliferation (CTV – CellTraceViolet) in CD8+ and CD4+ T cells after 6-day co-culture with stimulated DCs and **J**) Quantification. *k*= 6 co-cultures from *n*=5 donors across 2 experiments.

To further investigate the direct effects of iaDCs on T cell activation and polarisation, we identified all T cells within our dataset that neighboured either an iaDC or an aDC (Figure 4D, S9B). Direct comparison of iaDC-neighbouring and aDC-neighbouring T-cells revealed an upregulation of markers associated to T-cell activation (*IFNG*, *LAG3*, *GZMK*, *ICOS*) and effector memory differentiation (*SELL*, *SLAMF7*, *CD160*) to be enriched in T-cells directly adjacent to iaDC (Figure 4E, F). Whereas T-cells around aDCs showed expression of a classical activation profile, including *CD69*, *CTLA4*, *IL7R* and *IL2RA* (Figure 4E). Interestingly, this iaDC-adjacent signature was broadly induced across the T cell lineage, suggesting that iaDCs can orchestrate inflammation by directing diverse T cell populations (Figure 4G, S9C). To unravel the molecular relationship between the induction of EM CD4^+^ and CD8^+^ T-cells by iaDC, we isolated blood-derived DC2 from healthy donors, exposed them for 1 hour to either medium, IFNγ, or UPA, washed them, and cocultured them with naïve allogenic total T-cells for 6 days (Figure 4H). Exposure to UPA pretreatment reduced the capacity of DC2s to induce expression of IFNγ^+^ and proliferation in CD8^+^ and CD4^+^ T-cells (Figure 4I, J). On the other hand, pretreatment of DC2 with IFNγ significantly enhanced the induction expression of IFNγ^+^ and proliferation in CD8^+^ and CD4^+^ T-cells (Figure 4I, J). Of note IFNγ pretreatment of DC2 lead to an enhanced ability of iaDC to induce IFNγ expression in both CD4^+^ and CD8^+^ T-cells, thus revealing a direct feed-forward loop of iaDC function to T-cell phenotype. Hence, this data shows that similar to our *in vivo* findings IFNγ induced DC2 priming induces iaDC transcriptional priming and enhances generation of EM CD4^+^ and CD8^+^ T-cell phenotypes *in vitro*.

### The abundance of iaDC within inflamed areas of Crohn‘s disease-affected intestines predicts disease severity

To understand if induction of a chronic inflammation associated program in iaDC is specific to the lymph node or can be generalized to other chronic inflammatory diseases we analysed scRNAseq data from Crohn’s disease (CD) and ulcerative colitis (UC) patients, synovial membrane data of osteoarthritis (OA) and rheumatoid arthritis (RA) patients and of patients affected by skin sarcoidosis, and projected them onto our LN mononuclear phagocyte reference scRNAseq data (Figure 5A). Patients either presenting with an unresolving disease phenotype (GIMATS^High^) in CD (Figure 5B), or higher disease activity scores in UC, CD (Figure 5C, S10A), RA (Figure 5D, S10B) as well as in lesional areas of skin affected by sarcoidosis (Figure 5E, S10C) presented with significantly higher amounts of iaDCs present, indicating that iaDC associate with higher disease severity and may play a crucial role in regulating disease activity and progression.

**Figure 5.**
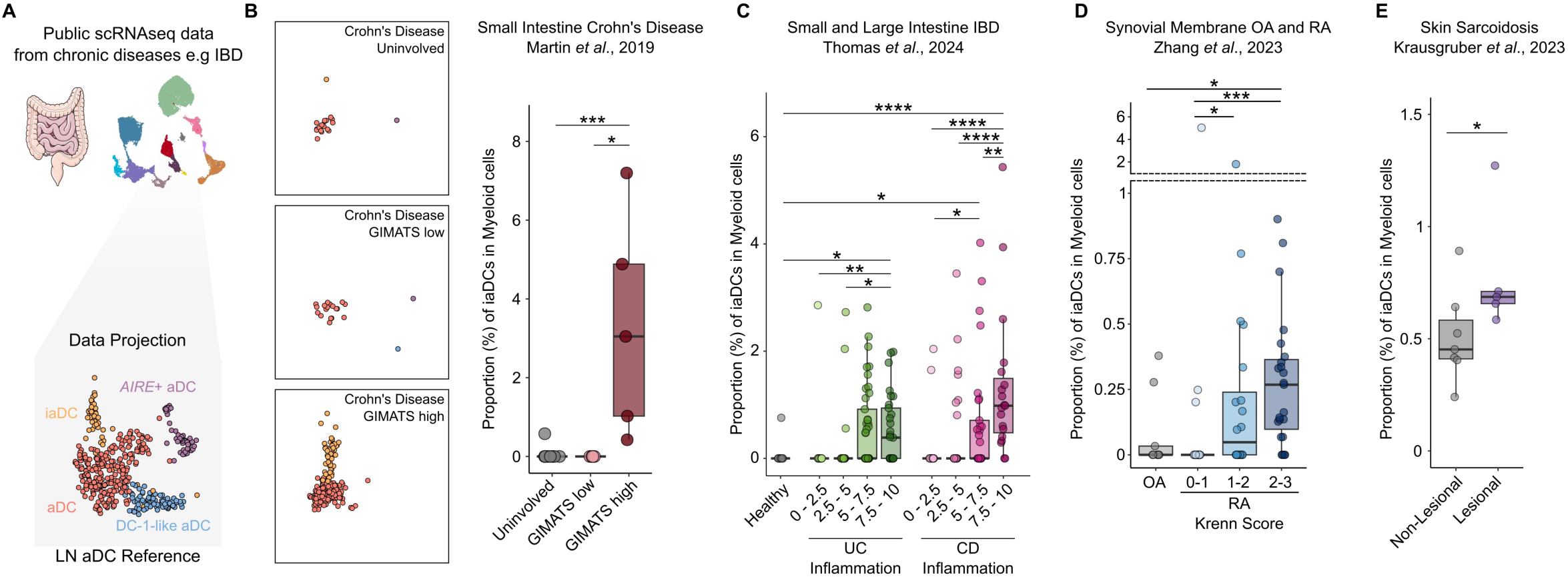
iaDC are a conserved feature of chronically inflamed human tissues. **A)** Schematic representation of data analysis for iaDC identification in public scRNAseq datasets. **B**) Projected UMAP embeddings of CCR7+ DCs from Small Intestinal Crohn’s Disease from (Martin 2019), stratified by “GIMATS” inflammation status and quantification of iaDC abundance by donor. **C**) Quantification of iaDC abundance in Ulcerative Colitis (UC) and Crohn’s Disease (CD) samples from (Thomas 2024), stratified by inflammation score. **D**) Quantification of iaDC abundance in Osteoarthritis (OA) and Rheumatoid Arthritis (RA) samples from (Zhang 2023) stratified by histopathological Krenn score. **E**) Quantification of iaDC abundance in non-Lesional and lesional skin biopsies from Sarcoidosis patients from (Krausgruber 2023).

### iaDCs localize to areas of active inflammation in Crohn’s disease and act as a spatial classifier of TNF blockade sensitivity

Presence and maintenance of effector EM CD4+ and CD8+ T-cells is a feature of chronic organ inflammation and has been linked to progression and resistance to treatment in Crohn’s disease. To mechanistically dissect the role of iaDC in Crohn’s disease and their link to higher disease activity scores, we analysed non-inflamed and inflamed regions of surgically resected small intestinal tissues from GIMATS^high^ patients using spatial transcriptomics (Figure 6A, S11A-E, Xenium, 10x Genomics). We identified a total of 77 clusters across the Myeloid, T/NK cell, B cell, Stromal, Endothelial and Epithelial cell lineages within 8 regions of interest (ROIs) (Figure S12). We found iaDCs to be present only within inflamed ROIs and with a strong local enrichment at the site of active inflammation (Figure 6B, C). Next, to better understand the composition and cellular relationships within the areas enriched for iaDCs we annotated 20 spatial niches with discrete cellular compositions across inflamed and non-inflamed ROIs with the dataset (Figure 6D, E, S13A, B). iaDCs were observed in Niche 0, 7, 8, 5 and 16 within our dataset, but were most strongly enriched in Niche 5. In Niche 0 and 7 iaDC co-enriched with EM memory and activated CD4+ and CD8+ T-cell phenotypes (*GZMK*+ CD8 T cell, *KLRB1*+ CD8 T cell, *TCF7*+ CD8 T cell and *KLRB1*+ CD4 T cell) and NK cells (*<u>CD160</u>*+ NK and *GNLY*+NK). Thus in conjunction with their microanatomical follicle like distribution around the Crohn’s disease induced ulceration we conclude that Niche 0 and 7 constitute tertiary lymphoid structures. On the other hand Niches 5 and 16, found at the leading edge of the inflammation-induced ulceration were strongly enriched for regulatory T-cells (Treg) with an unusal proinflammatory gene expression profile (Figure S13C), monocytes (*CXCL9*+ cMono; *SPP1*+ cMono), classically activated DCs (aDC) and activated Neutrophils (*CXCL8*+ Neut, *IL1B*+ Neut). Our prior data indicated a crucial role of IFNγ signalling in imprinting an iaDC transcriptional program on activated DCs (Figure 2,3,4). Here mapping of *IFNG* expression on the spatial level revealed that *IFNG* expression colocalized with iaDC enriched niches 5 and 16 at the leading ledge of the ulceration (Figure 6F). This observation was further supported by an enrichment of the JAK-STAT signalling pathway in iaDCs and spatial niches using PROGENy (Figure 6G, S13C). We observed tightly regulated transcriptional and cellular spatial architecture around CD ulcerations in our dataset, with iaDCs localised close to the borders of ulcerations (Figure 13E-H).

**Figure 6.**
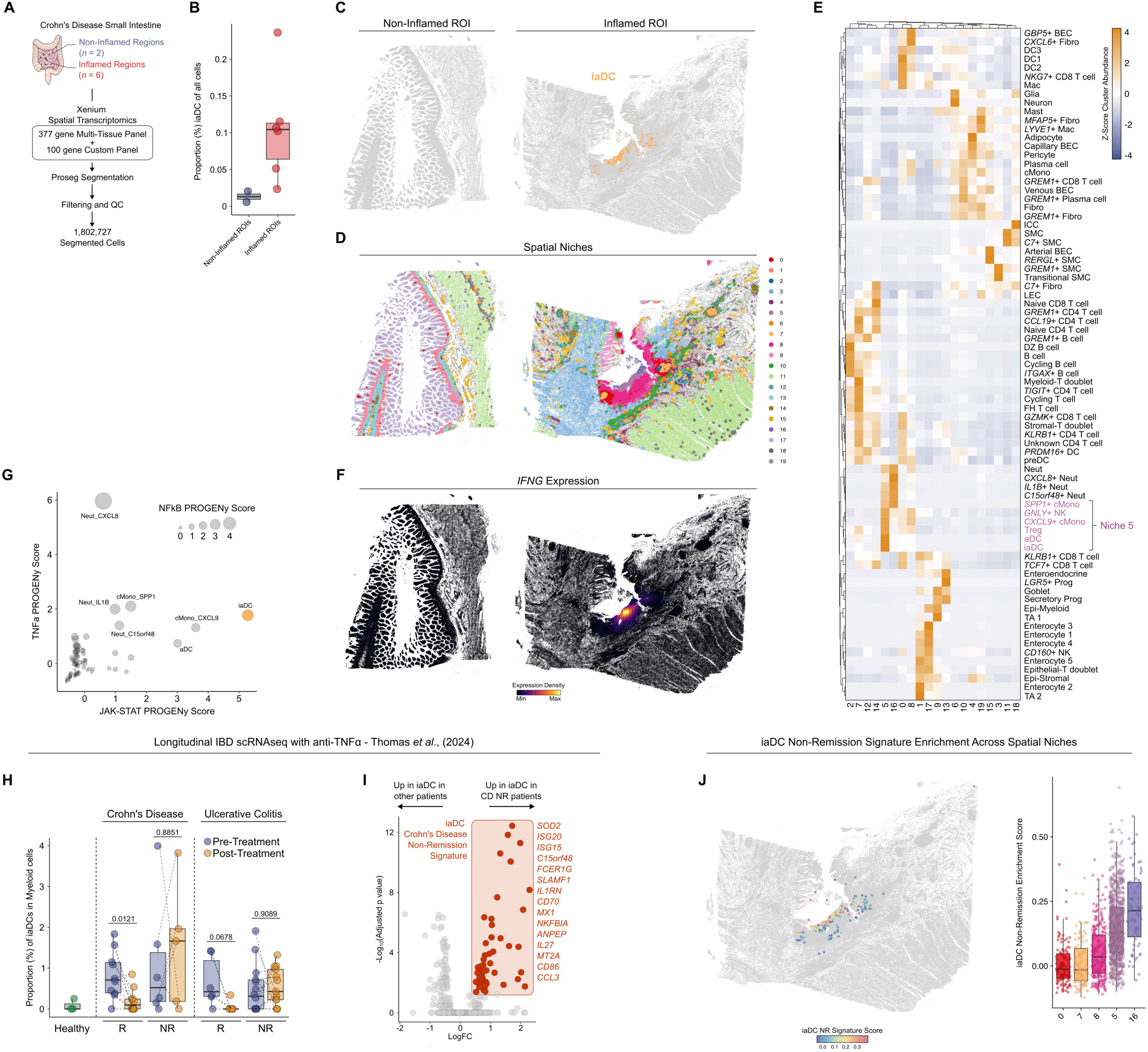
A spatially-restricted iaDC gene signature predicts response to anti-TNFa therapy. **A)** Schematic representation of Xenium experimental and pre-processing workflow. **B**) Quantification of iaDCs abundance across ROIs profiled. **C**). Visualisation of iaDCs across a representative non-inflammatory and inflammatory ROI. **D**) Visualisation of spatial niches across a representative non-inflammatory and inflammatory ROI. **E**) Heatmap depicting enrichment of Xenium clusters across calculated spatial niches. Clusters enriched in Niche 5 (iaDC-containing) are highlighted. **F**) Visualisation of *IFNG* expression across a representative non-inflammatory and inflammatory ROI. **G**) Scatterplot depicting JAK-STAT, TNFa and NFkB signalling scores across Xenium clusters, as determined via PROGENy analysis. **H**) Quantification of iaDC abundance in Crohn’s Disease (CD) and Ulcerative Colitis patients before and after anti-TNFa treatment (Thomas 2024), stratified by response status. R – Remission, NR – Non-Remission. **I**) Volcano plot of DEGs between iaDC from CD NR patients vs all other patients. iaDC CD NR signature genes are highlighted in red. **J**) Left, Visualisation of iaDC CD NR signature enrichment in Xenium iaDCs in an inflammatory ROI, and right, quantified across the total dataset, stratified by iaDC spatial niche localisation.

Resistance or refractoriness to anti-TNFα therapy is common in GIMATS^high^ patients. Thus, we aimed to determine whether the presence and intratissular localisation of iaDC can be utilized to predict therapy response. To do so, we first analysed the abundance of iaDCs within a recently published longitudinal single-cell transcriptional atlas of anti-TNFα treatment, examining pre- and post-treatment biopsies from anti-TNFα responders and non-responders affected by Crohn’s disease or ulcerative colitis (Figure 6H, S14A). Within this cohort, the response to anti-TNFα therapy was associated with a decline in the abundance of iaDC, whereas non-responders did not exhibit this decline in either disease. To mechanistically understand this relationship, we derived a non-remission iaDC signature by comparing iaDCs present in non-responders to all other iaDCs within this patient cohort and found that non-remission-associated iaDCs express *SOD2*, *ISG15*, *ISG20*, *C15orf48* and *SLAMF1* (Figure 6I, S14B-D). Next, we interrogated whether there is a specific spatial enrichment of the non-remission-associated iaDC signature across the spatial niches that harbouring iaDCs. Here we found that only clusters that localized to the edge of active inflammation within the ulceration (cluster 5, 16) were enriched for the iaDC non-remission transcriptional program (Figure 6J, S14E). Taken together, deconvolution of spatially restricted iaDC-associated non-remission signatures builds a spatial disease classifier to predict treatment efficacy.

### Cross-tissue and disease analysis reveals conserved iaDC neighbourhoods in human chronic inflammatory disease

Linking disease mechanisms across tissues and diseases can be beneficial to reveal common targets for drug development. Thus, to understand the transcriptional underpinnings of the observed association of iaDC presence with enhanced disease activity across various chronic inflammatory diseases, we integrated our LN data with available data from CD, UC, RA, and sarcoidosis patients, derived differentially expressed genes between classically activated DCs and iaDC for each dataset, and defined disease-specific and shared features across diseases (Figure 7A). This analysis revealed disease-specific genes mirroring their respective organ contexts, such as xxx for IBD, xxx for RA, and xx for Sarcoidosis (Figure 7B, C). Additionally, a conserved subset of genes across disease, independent of tissue origin, was detected (Figure 7D). Here many of the genes implicated in the mechanistic basis of iaDC effector memory generation were included, such as *CXCL9*, *CXCL10*, *CXCL11*, the IFNγ responsive gene *STAT1*, the antiapoptotic molecule *BCL2A1* and the effector and recognition genes *SLAMF7*, *CLEC7A*, *IDO1*, *C1QB*, *HLA-DMA* and *FCER1G*. To further understand whether the spatial niche orchestrated by this program is conserved across diseases, we generated and analysed spatial transcriptomics data from CD, sarcoidosis-affected lymph nodes and lungs, and RA synovial membranes (Figure 7E). After harmonisation of cell cluster annotation using a custom-trained Celltypist prediction model (Figure S15A-C), we analysed the distributions of iaDCs and the enrichment of the transcriptional signature of the chronic inflammatory niche identified in LNs onto all analysed samples (Figure 7F). This analysis showed that iaDCs clusters in specific micro-anatomical neighbourhoods across disease. In CD samples, as shown before, iaDC mainly concentrated to the ulceration area and the surrounding lymphoid follicles, which also coincided with the enrichment of the chronic inflammatory niche score (Figure 7F). Furthermore, in both sarcoidosis-associated LN and lung tissues, the presence of iaDCs (Figure 7F) exhibited the same pattern as the enrichment of the chronic inflammation-associated gene program, and a similar trend was observed within the RA-affected synovium (Figure 7F). To determine whether this cross-disease enrichment is based solely on the iaDC transcriptional signatures or on the conservation of a specific iaDC-driven cellular neighbourhood present across the analysed diseases, we performed a spatial connectivity graph-based neighbourhood enrichment analysis (Figure 7G). Through this analysis we identified the cellular components across tissues and diseases which were either statistically over-or under-represented directly adjacent to iaDCs *in situ*. Comparing cluster enrichment scores across all analysed tissues and diseases revealed a conserved iaDC cellular neighbourhood, enriched for effector memory T cells, *CXCL9+* Monocytes, conventionally aDCs, Tregs and *ITGAX*+ NK cells (Figure 7H, I). These results support the notion of iaDC and their associated cellular neighbourhood as defining feature of human chronic inflammatory diseases, such as IBD, sarcoidosis and RA.

**Figure 7.**
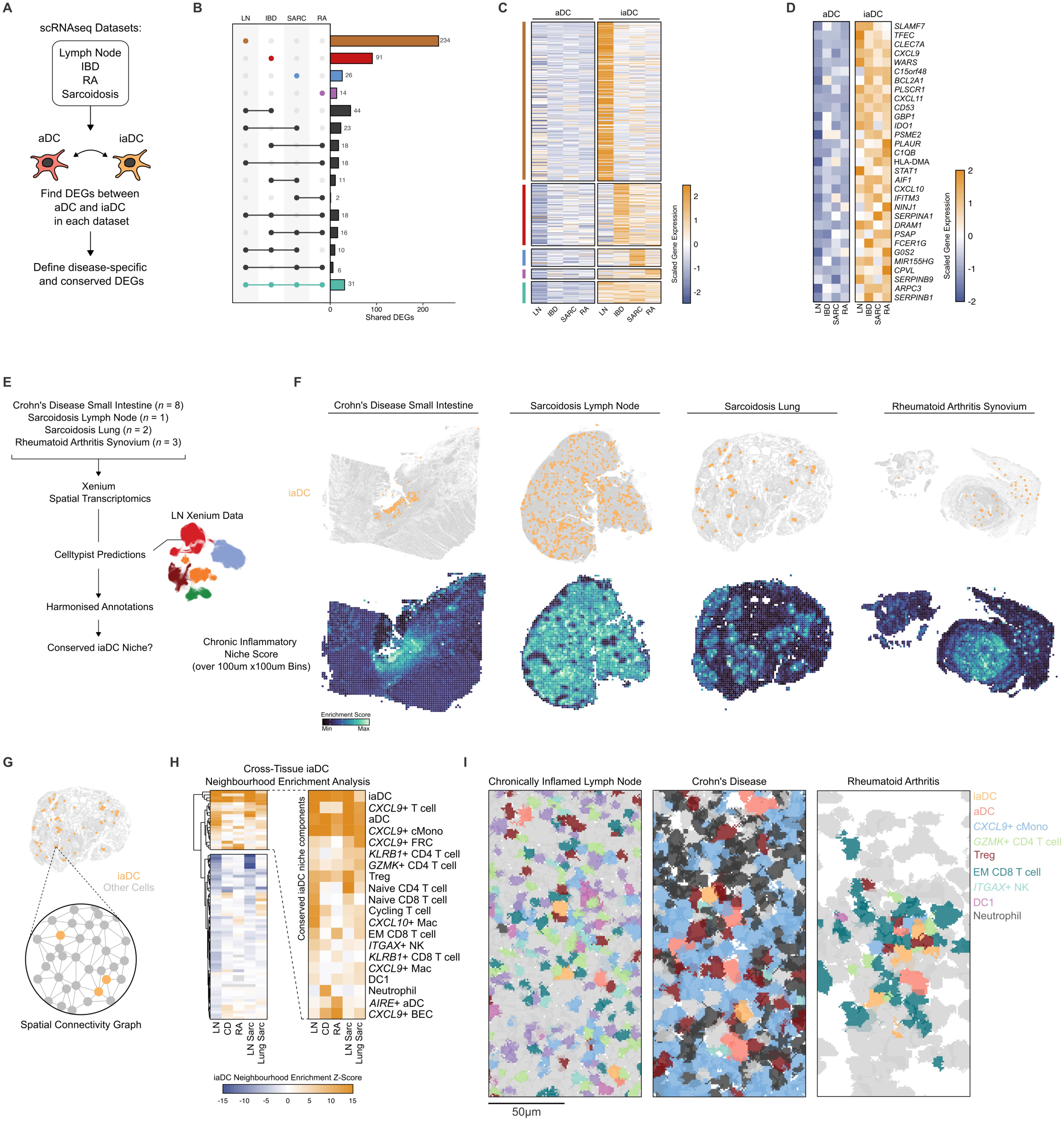
iaDC occupy a conserved niche in chronically inflamed human tissues. **A**) Schematic representation of data analysis for defining conserved and disease-specific features of iaDCs. **B**) Upset plot showing overlap between iaDC DEGs from different datasets, and **C**) heatmap depicting gene expression of divergent and conserved genes in aDCs and iaDCs from each dataset, as indicated in H. **D**) Heatmap depicting gene expression of the core iaDC gene signature in aDCs and iaDCs from each dataset. **E)** Schematic representation of Xenium experimental and Celltypist predictions workflow. **F**) Top, Visualisation of iaDCs across chronically inflamed human tissues profiled by Xeniium. Bottom, Visualisation of LN chronic inflammatory niche (Niche 5 – Fig2) gene signature enrichment scores in 100×100μm binned Xenium data from chronically inflamed human tissues. **G**) Schematic representation of the iaDC spatial connectivity graph for a lung sarcoidosis Xenium sample. **H**) Heatmap depicting enrichment scores for all clusters in local iaDC spatial neighbourhoods, across chronically inflamed human tissues. Zoomed section, cellular components of the conserved iaDC niche. **I**). Spatial plots depicting Proseg segmented cells within chronically inflamed human tissues, clusters in conserved niche are highlighted.

## Discussion

Chronic inflammatory diseases, such as Crohn’s disease, rheumatoid arthritis or sarcoidosis, pose a significant clinical challenge. Only limited and often non-specific treatment options exist; thus, the discovery of novel cellular and molecular disease mechanisms and drug targets remains of utmost importance. Progress in identifying novel therapeutic targets is often hindered by the heterogeneous nature of chronic inflammation, making it challenging to stratify disease phenotypes based on molecular and cellular patterns. Here, we show that human DC2 undergo a specific, chronic inflammation-associated transcriptional programming directed by IFN-γ-elicited JAK/STAT signalling within human cervical lymph nodes, Crohn’s disease- and ulcerative colitis-associated small intestine, and sarcoidosis-affected lungs and lymph nodes, entering the inflammatory activated DC (iaDC) cell state. Here, iaDCs are located in specific conserved cellular neighbourhoods, colocalizing with effector CD4+ and CD8+ EM T-cells, NK cells and effector-type myeloid cells, including *CXCL9*+ monocytes and DC1s. Across diseases, the presence of iaDC or their associated niches correlated with higher disease activity and refractoriness to treatment.

Prior studies have revealed that DC activation plays a pivotal role in shaping T-cell immunity ^7^. In particular both human and murine studies have shown that selective activation of DC subtypes DC1 and DC2, produces different outcomes in regards to functional T-cell polarization ^20,21^. However, these studies disregarded the intra-tissular signalling cues shaping DC functionality during ongoing inflammation, as found in cancer, acute or chronic inflammatory diseases. Upon activation DCs undergo tightly regulated transcriptional, phenotypic and functional reprogramming. In cancer this state has been previously termed “mature dendritic cells enriched in immunoregulatory molecules” (mregDC), characterized by their uniform expression of CCR7, LAMP3 and FSCN1 across multiple tumour entities ^10^. In other studies, an analogous state has been shown to be induced in DCs upon homeostatic uptake of apoptotic cells. Collectively, these cells can be referred to as activated DCs (aDCs). iaDCs, identified here, also express CCR7, LAMP3, FSCN1 and PD-L1, however, in contrast to aDCs, they additionally express high levels of pro-inflammatory, interferon-stimulated cytokines, such as CXCL9, CXCL10, CXCL11 and display high levels of JAK/STAT signalling. Furthermore, we show that their emergence is induced upon exposure to IFN-γ in local spatial niches, and not due to apoptotic cell debris uptake or sensing of other antigens ^8^. The distinct phenotypic and functional characteristics and developmental trajectory of iaDCs clearly demarcate them as a distinct transcriptional state of activated DCs. Furthermore cancer-associated and homeostatic aDCs have been largely associated with the DC1 lineage, whereas our data clearly support a DC2 ontogeny of iaDCs.

The exact cues which lead to the induction of aDCs within the tumour microenvironment and the tumour-draining lymph nodes are not known. However prior studies have provided candidates, such as AXL, TNFα, IFNγ and PGE_2_, which likely contribute to DC activation ^8,22^. During chronic inflammation we identify IFNγ as a crucial imprinter of the transcriptional identity of iaDC. Utilizing UPA, a clinically approved inhibitor of the JAK-STAT signalling pathway, we can show that spatially restricted IFNγ signalling by bystander cells, such as NK and T-cells is crucial to iaDC transcriptional identity. Interestingly IFNγ secretion was restricted to the immediate neighbours of iaDC in our spatial data, and was induced in naïve T cell by iaDCs *in vitro*. These results further highlight the unique spatial organisation necessary to fuel chronic inflammatory DC and T-cell organization, and point towards an IFNγ-dependent feed-forward loop mechanism which sustains chronic inflammation via an iaDC-T cell axis. Similar to aDC, iaDC and their transcriptional, phenotypic, functional and spatial features were conserved across several human tissues and chronic inflammatory conditions. Here core components of iaDCs were conserved, including their dependence on IFNγ and their spatial association with effector T-cell phenotypes *in situ*, allowing an association of iaDC presence and higher disease activity.

Collectively, we demonstrate that human chronic inflammation imposes a specific and spatially restricted transcriptomic program on activated DC2 within affected tissues. Furthermore, we show that iaDC transcriptional programming relies on cellular neighbourhood-specific IFN-γ signalling and leads to the preferential induction of effector memory CD4+ and CD8+ T-cell neighbourhoods within iaDC-dominated niches. Additionally, conserved functional, cellular, and transcriptional features of these iaDC effector niches are found across tissues and diseases, including Crohn’s disease, RA, and Sarcoidosis, and are associated with higher disease activity. This makes iaDCs a promising novel target to allow for the spatial stratification of chronic inflammatory diseases, especially in light of their susceptibility to JAK/STAT inhibition.

## Acknowledgments

This study was funded by the Deutsche Forschungsgemeinschaft (DFG, German Research Foundation) under Germany’s Excellence Strategy – EXC2151 – 390873048 (A.S., J.L.S), SFB 1454-P05-432325352 (A.S.) and University of Bonn TRA Research Prize.

## Supplemental Figure Legends

**Figure S1. Identification of CCR7+ DCs in human tissues via flow cytometry.** Gating strategies for the identification of CCR7+ DCs in human **A**) Lymph Nodes, **B**) Bronchioalveolar Lavage Fluid (BALF), **C**) Synovial Membrane, **D**) Synovial Fluid, **E**) Subcutaneous Adipose Tissue, and **F**) Peripheral Blood Mononuclear Cells (PBMCs).

**Figure S2. Profiling of human LN MNPs via scRNAseq and flow cytometry. A)** UMAP visualisation of scRNAseq of MNPs from human LNs, with level 1 (broad) clusters annotated. **B**) Dotplot depicting normalised expression of DEGs between broad clusters in scRNAseq data. **C)** UMAP visualisation of scRNAseq of MNPs from human LNs, with level 2 (fine) clusters annotated. **D**) Dotplot depicting normalised expression of DEGs between fine clusters in scRNAseq data. **E**) Quantification of CCR7+ DC subcluster abundances in scRNAseq data, stratified by sample type. **F**) Heatmap depicting Pearson’s Corelation of level 2 cluster abundances across samples profiled. **G)** Dotplot depicting normalised expression of marker genes encoding surface markers which discriminate iaDCs from aDC. **H**) Flow cytometry plots of CD274 (PD-L1) and CD319 (SLAMF7) within CCR7+ DCs in 2 human lymph node samples, depicting the variable abundance of CD274+CD319+ iaDC. **I**) Quantification of expression of makers identified in A, in CD274-CD319-aDC and CD274+CD319+ iaDC by flow cytometry.

**Figure S3 Construction of a human LN MNP scRNAseq atlas. A)** Table indicating the studies from which scRNAseq data was obtained, and the number of donors per disease group per study. **B**) UMAP visualisation of integrated LN MNP scRNAseq atlas, with cells derived from each study highlighted in navy on each panel. **C**) UMAP visualisation of integrated LN MNP scRNAseq atlas, with broad clusters annotated. **D**) Dotplot depicting normalised expression of DEGs between broad clusters in scRNAseq data. **E**) Subclustering of CCR7+ DCs within scRNAseq data. **F**) Overlay of previously identified iaDCs from in-house scRNAseq data (Fig 1B) onto integrated CCR7+ DC dataset. **G**) Dotplot depicting normalised expression of DEGs between CCR7+ DC subclisters. **H**) Quantification of iaDC abundance in integrated scRNAseq data stratified by disease group, and **I**) further stratified by disease sub-group in lymphoma and metastatic LNs.

**Figure S4 iaDC are induced by IFN**γ **and dependant on JAK-STAT signalling. A)** Violin plots depicting enrichment scores for response to Type II IFN, IFNα and IFNβ in level 2 (fine) clusters in scRNAseq data. **B**) Violin plots depicting scaled scores for JAK-STAT, TNFa and NFkB signalling in level 2 (fine) clusters in scRNAseq data, as determined by PROGENy analysis. **C**) Flow cytometry gating for the identification and export of LN MNPs after bulk stimulation cultures. **D**) Visualization of marker expression for all flow cytometry markers in LN MNPs after bulk stimulation cultures, overlain on UMAP embedding. **E**) Flow cytometry gating for the identification of CCR7+ DC2 in LN after bulk stimulation cultures. **F**) Expression and quantification of CXCL9, CXCL10, CD319 and CD274 in LN CCR7+ DC2s after stimulations. **G**) Flow cytometry gating for the isolation of DC2 from healthy PBMCs by FACS. **H**) Expression of CXCL9, CXCL10, CD274, CD319, CD273, CD86 and CD80 in Blood-derived DC2s after stimulations.

**Figure S5 Xenium Spatial Transcriptomics of human LNs. A)** UMAP visualisations of scRNAseq of MNPs from human LNs, left, UMAP computed on data downsampled to include only the 337 genes from Xenium predesigned Xenium Human Multi-Tissue and Cancer Panel. Right, UMAP computed on data downsampled to include only the 337 genes + 100 custom add-on genes. **B**) Quantification of prediction accuracy of LN MNP scRNAseq clusters (Dataset split in 2, half for training, half for testing, see Methods) when using predesigned panel only or predesigned + Custom add-on genes. *p > 0.05 **C**) UMAP visualisation of Xenium data from human LN, with cell lineages annotated, and **D**) individual UMAPs from each LN profiled grouped by disease status. **E**) Violin plot of number of genes expressed per Proseg segmented cell across LN samples. **F**) Visualisation and **G**) quantification of cell lineages across LN Xenium samples, grouped by disease status.

**Figure S6. Cluster Annotations in human LN Xenium Spatial Transcriptomics data. A)** Left, UMAP visualisation of LN Xenium Myeloid cells with subclusters annotated, Middle, Dotplot depicting normalised expression of DEGs between Myeloid cell subclusters in LN Xenium data. Right, Heatmap depicting relative abundances of Myeloid cell subclusters across LN samples. **B)** As in (A) but for T/NK cells. **C)** As in (A) but for B cells. **D)** As in (A) but for Stromal cells. **E)** As in (A) but for Endothelial cells.

**Figure S7. Spatial Niche analysis of human LN Xenium Spatial Transcriptomics data. A)** Schematic representation of spatial niche construction in Xenium data, a 50um raster scan is taken around every cell recording the identifying the cluster identities of all cells in the scanned area, and a cell-by-neighbour matrix is then computed across all cells, and cells are clustered by K-means clustering, to define spatial niches. **B**) Visualisation of spatial niches across LN Xenium samples. **C**) Quantification of spatial niches across LN Xenium samples, grouped by disease status. **D**) Multi-panel plot highlighting the localisation of each spatial niche and two niche-enriched clusters in a chronically inflamed LN profiled by Xenium. **E**) Visualisation of JAK-STAT signalling scores across 3 representative LN Xenium samples, as determined by PROGENy analysis. **F**) Heatmap depicting expression of *IFNG* in T/NK cell clusters from either Niche 1 and Niche 5 in LN Xenium data.

**Figure S8 Perturbation of human lymph nodes ex vivo with conserved spatial context. A)** UMAP visualisation of human LN slice culture Xenium data, with cell lineages annotated. **B**) Violin plot of number of genes expressed per Proseg segmented cell across LN slices. **C**) Dotplot depicting normalised expression of DEGs between cell lineages in LN slice culture Xenium data. **D**) Visualisation of cell lineages across LN Xenium slice samples. **E**) Heatmap of average JAK-STAT signalling scores across LN slice culture conditions, as determined by PROGENy analysis. **F**) Heatmap of average IFNγ-response signalling signature enrichment scores across LN slice culture conditions. **G**) Scatterplots depicting log-normalised average gene expression across LN slices from Xenium data. DEGs between slice conditions (adjusted p value <0.05, Wilcoxon rank sum test) are highlighted in red and blue. **H**) Visualisation of Celltypist-predicted iaDCs across LN Xenium slice samples. **I-K**) Heatmaps depicting **I**) relative abundances of Celltypist-predicted cluster identities across LN slice culture conditions, **J**) average IFNγ-response signalling signature enrichment scores within Celltypist-predicted clusters across LN slice culture conditions, and **K**) average JAK-STAT signalling scores within Celltypist-predicted clusters across LN slice culture conditions.

**Figure S9 Analysis of iaDC-T cell interactions in LN Xenium data. A)** Quantification of the distance from each aDC and iaDC to the nearest cell from each T/NK cluster across the Xenium dataset. **B**) Stacked barchart showing the total composition of aDC-neighbouring and iaDC-neighbouring T cells. **C**) Volcano plots of DEGs between all aDC-neighbouring and iaDC-neighbouring cells from each T cell cluster across the Xenium dataset. Significantly differentially expressed genes (logFC > 0.05 and < −0.05, adjusted p value <0.05, Wilcoxon rank sum test) are shown in red and blue.

**Figure S10 Conserved emergence of iaDCs in chronic inflammation in humans. A)** Projected UMAP embeddings of CCR7+ DCs from Ulcerative Colitis (UC) and Crohn’s Disease (CD) samples from (Thomas 2024), stratified by inflammation score. **B**) Projected UMAP embeddings of CCR7+ DCs from Osteoarthritis (OA) and Rheumatoid Arthritis (RA) samples from (Zhang 2023), stratified by histopathological Krenn score. **C**) Projected UMAP embeddings of CCR7+ DCs from non-Lesional and lesional skin biopsies from Sarcoidosis patients from (Krausgruber 2023).

**Figure S11 Xenium Spatial Transcriptomics of human Crohn’s Disease small intestine. A**) UMAP visualisation of Xenium data from human Crohn’s Disease small intestine (CD-SI) samples, with cell lineages annotated, and **B**) individual UMAPs from each CD-SI sample profiled grouped by inflammation status. **C**) Violin plot of number of genes expressed per Proseg segmented cell across CD-SI samples. **D**) Visualisation and **E**) quantification of cell lineages across CD-SI Xenium samples, grouped by inflammation status.

**Figure S12 Cluster annotations in human Crohn’s Disease small intestine Xenium data. A)** Left, UMAP visualisation of CD-SI Xenium Myeloid cells with subclusters annotated, Middle, Dotplot depicting normalised expression of DEGs between Myeloid cell subclusters in CD-SI Xenium data. Right, Heatmap depicting relative abundances of Myeloid cell subclusters across CD-SI samples. **B)** As in (A) but for CD4+ T cells. **C)** As in (A) but for CD8+ T cells. **D)** As in (A) but for B and Plasma cells. **E)** As in (A) but for Stromal cells. **F)** As in (A) but for Endothelial cells**. G)** As in (A) but for Epithelial cells.

**Figure S13 Spatial niche analysis of human Crohn’s Disease small intestine Xenium data. A)** Visualisation of spatial niches across CD-SI Xenium samples, grouped by inflammation status. **B**) Stacked barchart showing the total spatial niche composition for each CD-SI Xenium sample. **C**) Volcano plots of DEGs (logFC > 0.05 and < −0.05, adjusted p value <0.05, Wilcoxon rank sum test) between Tregs from Niche 5 (red) and all other Tregs (Blue). **D**) Scatterplot depicting JAK-STAT, TNFa and NFkB signalling scores across spatial niches, as determined via PROGENy analysis. **E)** Zoomed FOV of an ulceration from an inflamed CD-SI ROI. **F**) FOV with cell coloured by their distance to the ulceration border. **G**) Heatmap depicting gene expression of cytokines and chemokine gene expression, binned by distance (50um bins) from ulceration border. **H**) Left plots, Spatial plots of Xenium cluster with specific spatial patterning around CD ulcerations and Right plots, Quantification of cluster abundances per 50um bin, for 5 CD-SI samples which contain ulcerations.

**Figure S14 Analysis of iaDCs in Non-remission CD patients. A)** Projected UMAP embeddings of CCR7+ DCs from Ulcerative Colitis (UC) and Crohn’s Disease (CD) samples from (Thomas 2024), stratified by Treatment and Remission status. **B**) Heatmap of gene expression of iaDC CD NR signature genes in iaDCs from healthy patients, UC and CD patients, stratified by remission status. R – Remission, NR – Non-Remission. Genes overlapping with Xenium panel are highlighted in orange. **C**) Heatmap of iaDC CD NR signature enrichment scores across predicted clusters in scRNAseq data from (Thomas 2024), stratified by disease and remission status. **D**) Heatmap of iaDC CD NR signature enrichment scores across original cluster annotations from (Thomas 2024), stratified by disease and remission status. **E**) Left, Visualisation of spatial niches in an additional inflammatory CD-SI ROI. Right, Visualisation of iaDC CD NR signature enrichment in iaDCs in an additional inflammatory CD-SI ROI.

**Figure S15 Xenium cluster harmonisation using Celltypist. A)** Heatmap showing the proportion of Celltypist predicted annotations (x-axis) derived from each original manual clustering annotations (y-axis) from the CD-SI xenium dataset. **B**) Dotplot depicting normalised expression of DEGs between Celltypist predicted annotations CD-SI Xenium data. **C**) Violin plot of number of genes expressed per Proseg segmented cell across Lymph Node (LN) sarcoidosis, Lung sarcoidosis and Rheumatoid Arthritis samples.

## Materials and Methods

### Patient Samples and Ethics

Human tissue samples were obtained from patients who have given their written informed consent and were either undergoing surgical resection of cervical lymph nodes (after diagnosis of lymphadenopathy via ultrasound), Bronchioalveolar Lavage for diagnostic purposes, or ultrasound-guided synovial membrane and fluid sampling, at the University Hospital Bonn. All samples were obtained under approved ethical protocol of the University Hospital Bonn to. Dr. Thorsten Send and Prof. Dr. Andreas Schlitzer (Serial no. 251/20). **Add others**

### Tissue Processing

All samples were transferred from the University Hospital Bonn to the lab on ice and processed immediately upon receipt.

#### Lymph Nodes

LNs were washed in PBS (PAN-Biotech) to remove any blood, and any blood clots, fat, connective tissue or black cauterized tissue was cut away. LNs were cut into pieces using a scalpel before and finely minced (1mm^3^) using surgical scissors. LN fragments were then subjected to digestion with 150ug/ml Liberase TL low (Roche) and 100ug/ml DNAse I (Roche) in RPMI (PAN-Biotech) for 45-60mins with agitation at 37°C. The digestion was halted by the addition of RPMI supplemented with 10% Fetal Calf Serum (FCS) (Sigma-Aldrich), and the digested cell suspension was passed through a 100μm filter, and washed through with RPMI+10% FCS. Digested cell suspensions were pelleted, washed in FACS buffer (PBS + 2% FCS + 2mM EDTA (Invitrogen)) and counted for downstream use.

#### BALF

BALF samples were aspirated up-and-down with a stripette to break up any mucus clumps, and then passed through a 100μm filter. The same filter was then washed through with an additional 50ml of FACS buffer to collect any additional cells. Cell suspensions were pelleted, pooled and subjected to red blood cell (RBC) lysis (4mins at RT) with RBC-lysis Bffer (Biolegend) before washing with FACS buffer. Samples were re-filtered through a 100μm filter and counted for downstream use.

#### Synovial Membrane

Synovial biopsies were washed in PBS and finely minced (1mm^3^) using surgical scissors before digestion with 150ug/ml Liberase TL low and 100ug/ml DNAse I in RPMI for 45-60mins with agitation at 37°C. The digestion was halted by the addition of RPMI supplemented with 10% FCS, and the digested cell suspension was passed through a 100μm filter, and washed through with RPMI+10% FCS. Digested cell suspensions were pelleted, washed in FACS buffer (PBS + 2% FCS + 2mM EDTA) and counted for downstream use.

#### Synovial Fluid

Synovial fluid was diluted at least 5-fold in FACS buffer to account for viscosity and pelleted, before washing with FACS buffer. Cell suspensions were pelleted and subjected to RBC lysis (4mins at RT) before washing with FACS buffer. Samples were re-filtered through a 100μm filter and counted for downstream use.

#### Adipose Tissue

Adipose Tissue samples were cut into pieces using a scalpel before and finely minced (1mm^3^) using surgical scissors. Samples were digested with 200ug/ml Collagenase IV (Sigma-Aldrich) and 100ug/ml DNAse I in HBSS (PAN-Biotech) for 60mins with agitation at 37°C. The digestion was halted by the addition of RPMI supplemented with 10% FCS, and the digested cell suspension was passed through a 100μm filter, and washed through with RPMI+10% FCS. Cell suspensions were pelleted and subjected to RBC lysis (4mins at RT) before washing with FACS buffer. Samples were re-filtered through a 100μm filter and counted for downstream use.

#### PBMC

Blood samples were diluted 1:1 in PBS and layered onto a Pancoll gradient (PAN-Biotech), and spun for 25 mins without break at 750g. The leukocyte layer was collected and washed with FACS until the supernatant was clear. Samples were then filtered through a 100μm filter and counted for downstream use.

### Flow Cytometry Staining and Data Analysis

Cell suspensions were stained for viability with 1:1000 BD Horizon Fixable Viability Stain 510 (BD Biosciences) for 20 mins at 4°C, and washed with PBS. Cells were blocked in human blocking buffer (5% human serum (Sigma-Aldrich), 1% rat serum (Sigma-Aldrich), 1% mouse serum (Sigma-Aldrich), 5% FCS and 2mM EDTA) for 15 mins prior to antibody staining. For certain markers (CCR7, CADM1, CD127) antibody staining was performed prior to other antibodies, at 37°C for 25 mins. Cells were then washed and resuspended in FACS buffer and stained with the remaining antibody cocktails at 4°C for 30 mins. Cells were washed in FACS buffer and, if sorting, were resuspended directly in FACS buffer, and if phenotyping, were fixed for 20 mins at 4°C in Cytofix (BD Biosciences), before resuspension in FACS buffer. For intracellular staining, cells were fixed and permeabilised with BD Cytofix/Cytoperm Kit (BD biosciences), according to manufacturer’s instructions, and stained with antibodies specific for intracellular antigens for an additional 30 mins at 4°C. Flow cytometry was performed using a SONY ID7000 spectral analyser (SONY) or cells were purified by cell-sorting using a BD FACS Aria III (BD Biosciences). Flow cytometry data was analysed using FlowJo v10.8.1 (Treestar), R version 4.2.1 (The R foundation) and Prism 9 (GraphPad).

Downsampled Gated flow cytometry data was exported from flowjo, and imported into R and convertd into a Seurat object. Flow datasets were batch corrected using the ‘*RunFastMNN’* function in the SeuratWrappers R package, correcting for the sample of origin, before louvian clustering and UMAP visulaisation.

### LN digest stimulation assays

Cryopreserved (Cryostor CS10, Stemcell Technologies) LN cells were thawed, washed and resuspended in RPMI with 10% FCS and 1% Penicillin/Streptomycin (Pen/Strep) /Gibco). LN cells were plated at a density of 2×10^6 cells per well in 1ml in a 24 well plate and were cultured for 19 hours at 37°C 5% C0_2_ with or without stimulation with either 50ng/ml IFNγ (Peprotech) or 1μM Upadacitinib (Merck). 1:1000 Golgiplug (BD Biosciences) and 1:1000 Glogistop (BD Biosciences) were added for the final 4 hours of culture. Cells were harvested from plates and analysed by intracellular flow cytometry as detailed above.

### Blood DC2 sorts and stimulations

PBMCs were obtained from healthy donor buffy coat samples as detailed above. Prior to antibody staining and sorting, cells were first depleted of lymphoid cells via incubation with biotin-labelled antibodies (CD3, CD19, CD20, CD56, CD7) and Streptavidin MicroBeads (Miltenyi Biotec). Cell suspensions were then passed through a LS Column (Miltenyi Biotec) in a magnetic field, and the flow-through collected. The resultant suspension was stained and sorted for DC2s as detailed above. After sorting, DCs were resuspended in RPMI with 10% FCS and 1% Pen/Strep and plated at a density of 5×10^3 cells per well in 200ul in a 96-well U-bottom plate. Cells were cultured for 18 hours at 37°C 5% C0_2_ with or without stimulation with either 50ng/ml IFNγ or 1μM Upadacitinib, and 1:1000 Golgiplug and 1:1000 Glogistop were added for the final 4 hours of culture. Cells were harvested from plates and analysed by intracellular flow cytometry as detailed above.

### Allogenic T cell stimulations

Naïve T cells were isolated from PBMCs from healthy donor buffy coat samples using the Pan-Naïve T cell isolation kit (Miltenyi Biotec). Resultant cells were then stained and sorted by FACS to further enrich for naïve cells (CCR7+CD45RA+) to ensure a pure population for stimulations. DC2s were isolated from the same buffy coat samples as detailed above. After sorting, DCs were resuspended in RPMI with 10% FCS and 1% Pen/Strep and plated at a density of 1×10^4 cells per well in 200ul in a 96-well U-bottom plate. DCs were cultured for 1 hours at 37°C 5% C0_2_ with or without stimulation with 50ng/ml IFNγ or 1μM Upadacitinib, and then co-cultured with 4×10^4 allogenic CellTraceViolet (Invitrogen) −labelled Naïve T cells for 6 days in 200ul RPMI with 10% FCS and 1% Pen/Strep. On day 6, cultures were stimulated with 50ng/ml phorbol-12-myristate 13-acetate (PMA) and 1ug/ml Ionomycin (Cell Activation Cocktail, Biolegend), and incubated for 5 hours, with 1:1000 Golgiplug and 1:1000 Glogistop were added for the final 4 hours of culture. Cells were harvested from plates and analysed by intracellular flow cytometry as detailed above.

### *Ex vivo* LN Slice Culture assays

LN samples were cleaned of any blood clots, fat, connective tissue or black cauterized tissue and bathed in 0.01% Digitonin for 1 second before being washed in PBS. LNs were secured in place with cyanoacrylate glue and slices were cut using a VT1200S Vibratome (Leica) with a velocity of 0.12mm/s and an amplitude of 1mm). Slices were transferred to Millicell cell culture inserts (Merck) in 6 well plates containing RPMI supplemented with 10% FCS, 1x Glutamax (Gibco), 1% Pen/Strep, 50uM b-mercaptoethanol (Gibco), 1mM sodium pyruvate (Gibco), 1x non-essential amino acids (Gibco) and 20mM HEPES (PAN Biotech). Slices were cultured for 24 hours at 37°C 5% CO_2_ with or without stimulation with either 50ng/ml IFNγ or 1μM Upadacitinib. Slices were fixed in 4% Paraformaldehyde (PFA) (Science Services) for 2 hours at 4°C, before dehydration and embedding in Paraffin.

### scRNAseq of LN MNPs

LN cells were enriched for MNPs via magnetic-based depletion of lymphoid cells by incubation with biotinylated CD3, CD19, CD20, CD7 and CD34 antibodies and then Mojosort Streptavidin Nanobeads (Biolegend) for 15mins at 4°C, before separation in a magnetic field. MNP-enriched cell suspensions were washed, counted and 16,000 cells were loaded for Next GEM Single cell 3’ scRNAseq (10x Genomics) according to manufacturer’s instructions, for a targeted recovery of 10,000 cells per sample. scRNAseq reactions and downstream library construction were performed according to manufacturer’s instructions. Libraries were then sequenced using an Illumina Novaseq (Illumina) with 150-bp paired-end reads.

### scRNAseq data analysis

scRNAseq data was processed with the CellRanger Single-Cell Software Suite (10x Genomics) to align reads to the GRCh38 human reference genome and to construct count matrices for each sample. Single cell count matrices were analysed in R version 4.2.1 (The R foundation) using Seurat v4.3.0. Cells with <1500 genes detected and >10% mitochondrial reads were deemed to be low quality and removed from downstream analyses. Data were then log-normalised using the *<u>NormalizeData</u>* function in Seurat and integrated following the seurat v4 integration workflow. Clusters were identified on the integrated data using the *FindNeighbors* and *FindClusters* functions in Seurat, and UMAP dimensionality reduction was performed using the *RunUMAP* function in Seurat, with default parameters. Cell lineages were annotated on the basis of expression of known marker genes and MNPs were sub-clustered and subjected to another round of integration, clustering and UMAP visualisation. MNP level 1 annotations were achieved with default clustering and detailed level 2 annotations were determined using the *FindSubCluster* function in Seurat. Differentially expressed genes (DEGs) between clusters were calculated using the *FindMarkers* function in Seurat and using a tf-idf based metric, and annotation of clusters into cell states was performed by inspection of DEGs for each cluster.

Gene signatures for DC-1 and DC-2 derived activated DCs were obtained from (Cheng 2021), and gene signatures for Response to Type II Interferon, Intereferon Beta and Interferon Alpha were retrieved from MsigDB using the *msigdbr* R package. Single-cell gene signature enrichment scores were calculated for each gene signature using the *AddModuleScore* function in Seurat. PROGENy signalling pathway scores were calculated per cell using the *progeny* function in the progeny R package.

For Nichenet analysis, we first defined an iaDC gene signature, by calculating DEGs between iaDCs and aDCs, and taking upregulated genes in iaDCs with log2FC > 0.5 and adjusted p values < 0.01. All genes expressed (non-zero expression in at least 10% of cells) in aDCs were used as the background gene expression for the analysis. We did not exclude potential ligands based on expression metric cut-offs in our dataset, since our analysis was limited to MNPs, which may not express some ligands of interest which would be expressed by other cell lineages. Thus, through this analysis we calculated a list of “hypothetical predicted ligands” based only on the transcriptional features of iaDCs and aDC. NicheNet analysis was completed using the default workflow and parameters, and top iaDC-regulating ligands were visualised using pheatmap.

### LN MNP scRNAseq Atlas

Publicly available scRNAseq data from healthy, lymphoma, reactive and metastatic LNs as well as tonsilitis tonsils were downloaded and loaded into R directly or converted from h5ad format using the *readH5AD* function in the zellkonverter R package. Each dataset was processed individually, as detailed above, and MNPs were identified and extracted from each. MNPs from all datasets, and data from this study, were then merged and integrated following the seurat v4 integration workflow. Clusters were identified on the integrated data using the *FindNeighbors* and *FindClusters* functions in Seurat, and UMAP dimensionality reduction was performed using the *RunUMAP* function in Seurat, with default parameters. The CCR7+ DC cluster was re-clustered using the *FindSubCluster* function in Seurat, to identify putative iaDCs, aDCs and *AIRE*+ aDC.

### Public scRNAseq data projection

Publicly available scRNAseq data from Crohn’s Disease, Ulcerative Colitis, Osteoarthritis, Rheumatoid Arthritis and Sarcoidosis skin samples were downloaded and loaded into R directly or converted from h5ad format using the *readH5AD* function in the zellkonverter R package. MNPs were identified and extracted from each dataset, either by using publication annotations or via processing and clustering as detailed above. For each dataset, using the LN MNP data as a reference, we used the Seurat *FindTransferAnchors* function to find anchors between the reference and query datasets, and the *MapQuery* function to transfer cell labels from the reference to the query dataset, and to project query cells into the reference UMAP embedding. The abundance of iaDCs in query MNP datasets was then quantified.

### Xenium Spatial Transcriptomics – Panel design

For Xenium Spatial Transcriptomics experiments we used the Xenium Multi-Tissue and Cancer panel (377 genes), supplemented with 100 custom add-on genes. Custom genes were selected by defining uniquely-expressed genes for each cluster using a tf-idf based metric and by the inclusion of additional marker and functional genes from the literature. Proposed custom genes were then benchmarked for their ability to identify scRNAseq MNP clusters using a prediction-based approach. Briefly, LN MNP scRNAseq data was split randomly in half, into training and test datasets. The datasets were downsampled to include only the 377+100 genes in the proposed total panel, and the cell identities in the test dataset were predicted using the training dataset as a reference, using the *FindTransferAnchors* and *TransferData* functions in Seurat. The predicted annotations in the test dataset were then compared against the original ground-truth whole transcriptome annotations to assess accuracy of prediction and therefore the suitability of the panel to resolve the different populations. This process was iterated, with changes to the custom panel, in order to optimise panel design. This resulted in a final custom panel with 46 MNP identity/state defining genes, 19 identity/state defining genes of other lineages (T cells, B cells etc) and 35 functional genes.

### Xenium Spatial Transcriptomics – Workflow

Tissue sections were sectioned from FFPE blocks onto Xenium slides, followed by deparaffinisation and permeabilization following the 10x Genomics user guide CG000580. Probe hybridisation, ligation and amplification were then carried out on slides according to the instructions in 10x Genomics user guide CG000582. Slides were then loaded into the Xenium Analyser and acquired.

### Xenium Spatial Transcriptomics – Data Processing, Clustering and Visualisation

Output Xenium spatial transcriptomics datasets were re-segmented with Proseg using default parameters. Proseg-output count matrices were analysed in python using Scanpy and Squidpy. Cells with <15 counts per cell were removed from downstream analyses. Data from each sample per dataset were merged and spatial coordinates transformed so that all samples could be visualised on the same spatial embedding. Datasets were log-normalised using the *pp.normalize_total* and *pp.log1p* functions, and pca was performed using all genes in the datasets. The datasets were then integrated using the *harmony_integrate* function from the pyharmony package, adjusting principal components to correct for the sample of origin. The neighbourhood graph was then contructed on harmony-adjusted pca-components with the *pp.neighbors* function, and clustering and UMAP dimensionality reduction were then calculated with the *tl.leiden* and *tl.umap* functions respectively. Cell lineages were annotated on the basis of expression of known marker genes. Each lineage (B cell, T/NK cell, Stromal, Myeloid, Endothelial etc) was then sub-clustered and subjected to another round of integration, clustering and UMAP visualisation in scanpy. Differentially expressed genes (DEGs) between clusters within each lineage were calculated using the *tl.rank_genes_groups* function in Scanpy and using a tf-idf based metric, and annotation of clusters into cell states was performed by inspection of DEGs for each cluster and comparison of gene expression of known gene expression patterns in scRNAseq data.

For certain analyses and for visualisation Xenium datasets were loaded into R using the the *readH5AD* function in the zellkonverter R package. Xenium gene signature enrichment scores were calculated for each gene signature using the *AddModuleScore* function in Seurat, with modified paramters *ctrl* = 25, *nbin* = 10 to account for limited feature number in the dataset. PROGENy signalling pathway scores in Xenium datasets were calculated per cell using the *progeny* function in the progeny R package.

### Xenium Spatial Niche Analysis

In order to construct spatial niches in Xenium spatial transcriptomics datasets informed by the local cluster composition of the tissue we adopted a 50um-raster scan-based approach. For each cell in the dataset, a 50um radius was defined around the cell centroid, and the annotations of cells within the defined area were counted to construct a cell-by-neighbour matrix for each cell in the dataset. This matrix was log-normalised and scaled, before being clustererd by Kmeans clustering, using the *KMeans* function in the sklearn.cluster package. We performed this clustering over an iteration of k for each dataset, and a suitable value of k was selected for each dataset after manual inspection of the niches and the enrichment of clusters across niches.

### Xenium Cell Segmentation plotting

Proseg segmentation output *geojson* files were read into R using the *read_sf* function in the sf R package. Cell annotations and gene expression data from processed Xenium datasets were mapped to segmentation objects using the cell ID as a reference, and plotted on the segmented data using the *geom_sf* function in the ggplot2 package.

### Xenium Distance calculations

Distances between cell of interest in Xenium data were calculated using the *calc.shortest_distances_pairwise* function in the Monkeybread python package, selecting for the combinations of cell clusters of interest. For distances between aDC and iaDC and *IFNG*+ cells, *IFNG*+ cells were first defined as any cell in each dataset with non-zero expression of *IFNG,* and distance metrics were calculated in the same fashion.

### Xenium Neighbouring T cells Analysis

In order to determine the distinct features of iaDC and aDC-neighbouring T cells a spatial connectivity graph was first constructed between cells in the LN Xenium dataset using the *gr.spatial_neighbors* in the Squidpy python package. Next, the indices of iaDC, aDC and all T cell clusters were extracted from the spatial connectivity graph, and any T cells which were directly neighbouring iaDCs and aDCs in the graph were identified. These cells where then labelled with a new metadata column, which was subsequently used for visualisation and calculation of DEGs.

### Xenium Celltypist Analysis

In order to harmonise cell labels between our analyses of distinct Xenium datasets from different tissues and experiments a reference-based prediction approach using CellTypist was employed. First the LN Xenium dataset was subset to include only one chronically inflamed sample, which was deemed representative of the entire dataset due to the presence of all cell clusters and niches. The dataset was re-normalised to 10,000 counts per cell, to be consistent with the data formatting required for CellTypist analysis. A custom CellTypist model was then trained on the dataset using the *celltypist.train* function in the celltypist python package. Query Xenium datasets (LN slice models, Crohn’s Disease Small Intestine, Rheumatoid Arthritis Synovial Membrane and Sarcoidosis Lymph Node and Lung) were similarly re-normalised to 10,000 counts per cell and annotations were predicted using the *celltypist.annotate* function in the celltypist python package. Predictions were validated via comparison with original annotations and via examination of the gene expression profiles of predicted cells relative to the reference LN Xenium dataset.

### Binning Enrichment analysis

The presence of an analogous niche across human chronic inflammatory diseases was assessed by calculating and plotting the enrichment scores of a gene signature derived from the DEGs significantly upregulated in the LN Chronic Inflammatory Niche (Fig.3, Niche 5). Since this signature comprises, genes derived from heterogenous components of the niche, instead of calculating enrichment scores across single cells in these Xenium datasets, enrichment scores were instead calculated across 100×100μm binned Xenium data. 100×100μm bins were drawn over Xenium datasets, and transcript expression data from all cells with centroids within a given bin were averaged, to create a new binned dataset. These binned datasets were then tested for their enrichment of the Chronic Inflammatory Niche gene signature using the *tl.score_genes* function in Scanpy.

### Xenium iaDC neighbourhood enrichment Analysis

Spatial connectivity graphs were constructed for Xenium datasets with celltypist-harmonised annotations using the *gr.spatial_neighbors* function in the Squidpy python package. The enrichment of cell types within the spatial neighbourhoods of other cell types was then calculated with the *gr.nhood_enrichment* in the Squidpy python package. Enrichment Z-scores of cell types within iaDC-neighbourhoods in each dataset were then extracted and plotted with pheatmap.

### Data and Code Availability

Single cell and spatial transcriptomics data (count matrices and processed (.RDS and .h5ad files) will be made available on GEO upon publication of the final manuscript. All code required for the analysis of data will be available on Github upon publication of the final manuscript.

