## Supplemental Figures for "Human chronic inflammation is orchestrated by spatially restricted inflammation-activated Dendritic cells"

**Figure S1**

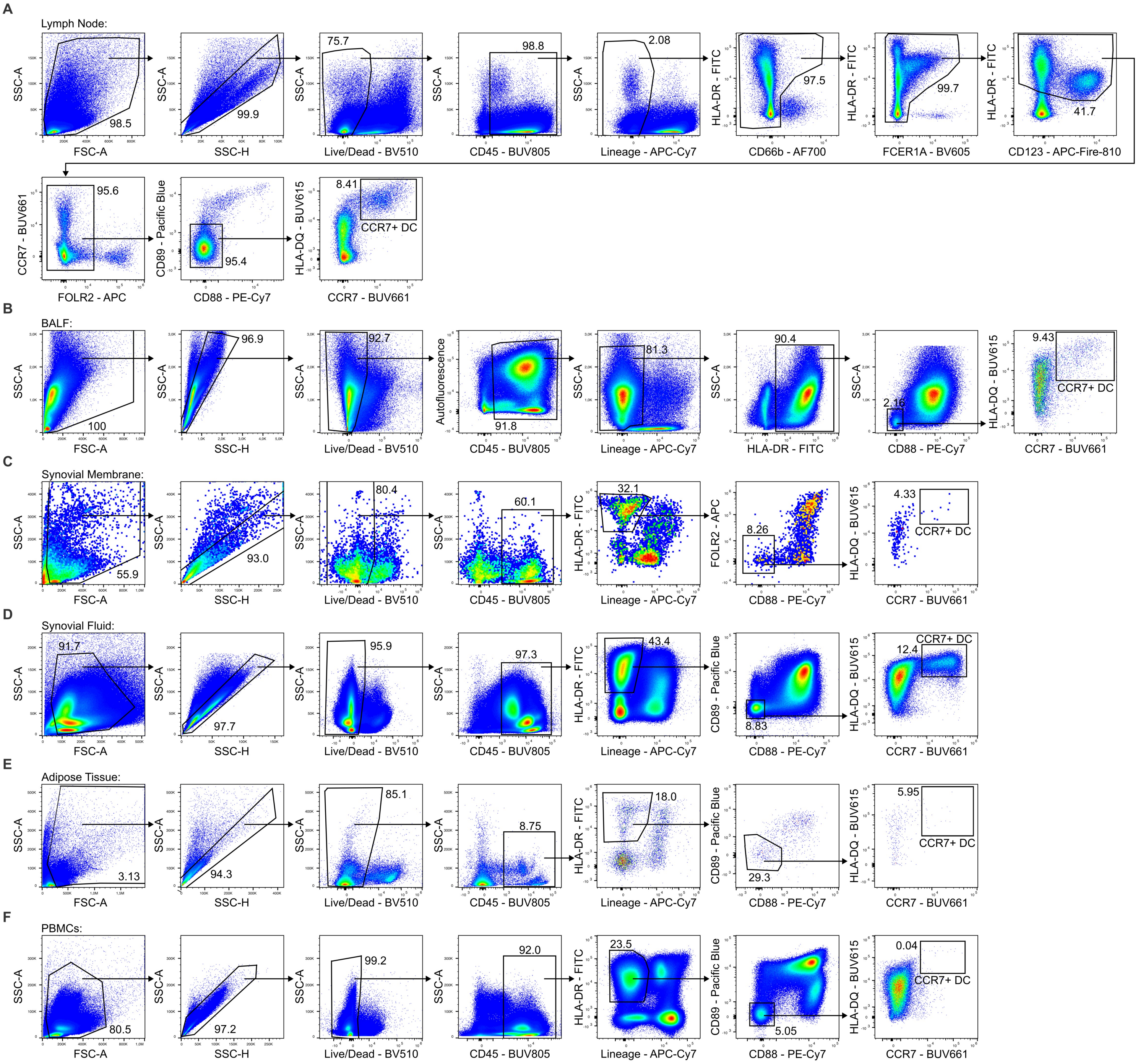
A
Lymph Node:
SSC-A
FSC-A
98.5
SSC-H
99.9
75.7
Live/Dead - BV510
98.8
CD45 - BUV805
2.08
Lineage - APC-Cy7
HLA-DR - FITC
97.5
CD66b - AF700
99.7
FCER1A - BV605
41.7
CD123 - APC-Fire-810
CCR7 - BUV661
95.6
FOLR2 - APC
CD89 - Pacific Blue
95.4
CD88 - PE-Cy7
HLA-DQ - BUV615
8.41
CCR7+ DC
B
BALF:
100
96.9
92.7
Autofluorescence
91.8
81.3
90.4
2.16
9.43
C
Synovial Membrane:
55.9
93.0
80.4
60.1
32.1
FOLR2 - APC
8.26
4.33
D
Synovial Fluid:
91.7
97.7
95.9
97.3
43.4
8.83
12.4
E
Adipose Tissue:
3.13
94.3
85.1
8.75
18.0
29.3
5.95
F
PBMCs:
80.5
97.2
99.2
92.0
23.5
5.05
0.04

Figure S2

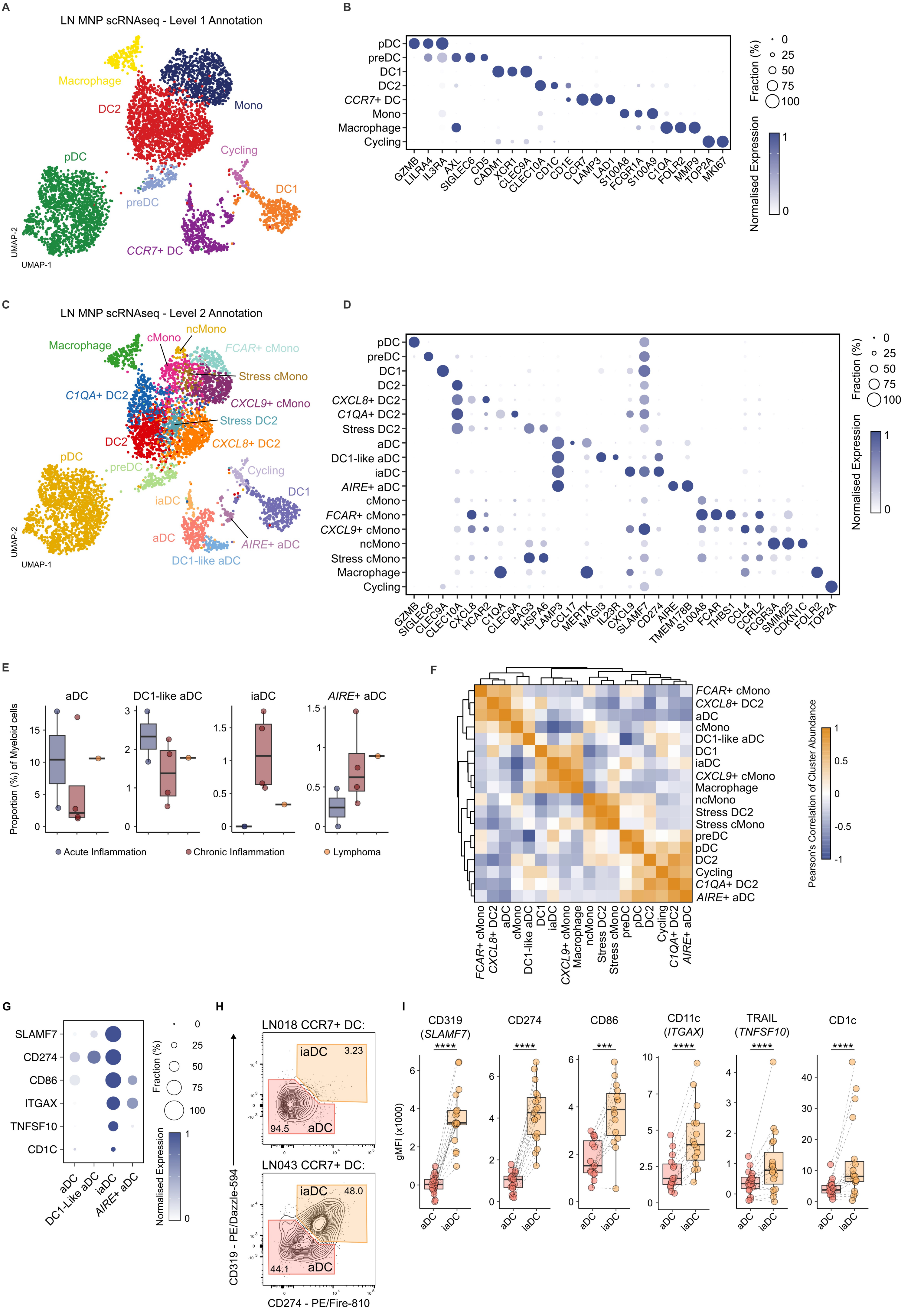

A
LN MNP scRNAseq - Level 1 Annotation
Macrophage
Mono
DC2
pDC
Cycling
preDC
DC1
CCR7+ DC
UMAP-2
UMAP-1
B
pDC
preDC
DC1
DC2
CCR7+ DC
Mono
Macrophage
Cycling
GZMB LILRA4 IL3RA AXL SIGLEC6 CD5 CADM1 XCR1 CLEC9A CLEC10A CD1C CD1E CCR7 LAMP3 LAD1 S100A8 FCGR1A S100A9 C1QA FOLR2 MMP9 TOP2A MKI67
Fraction (%)
Normalised Expression
C
LN MNP scRNAseq - Level 2 Annotation
Macrophage
cMono
ncMono
FCAR+ cMono
Stress cMono
C1QA+ DC2
CXCL9+ cMono
Stress DC2
DC2
CXCL8+ DC2
pDC
preDC
Cycling
DC1
iaDC
aDC
AIRE+ aDC
DC1-like aDC
D
pDC
preDC
DC1
DC2
CXCL8+ DC2
C1QA+ DC2
Stress DC2
aDC
DC1-like aDC
iaDC
AIRE+ aDC
cMono
FCAR+ cMono
CXCL9+ cMono
ncMono
Stress cMono
Macrophage
Cycling
GZMB SIGLEC6 CLEC9A CLEC10A CXCL8 HCAR2 C1QA CLEC6A BAG3 HSPA6 LAMP3 CCL17 MERTK MAGI3 IL23R CXCL9 SLAMF7 CD274 AIRE TMEM178B S100A8 FCAR THBS1 CCL4 CCRL2 FCGR3A SMIM25 CDKN1C FOLR2 TOP2A
Fraction (%)
Normalised Expression
E
aDC
DC1-like aDC
iaDC
AIRE+ aDC
Proportion (%) of Myeloid cells
Acute Inflammation
Chronic Inflammation
Lymphoma
F
FCAR+ cMono
CXCL8+ DC2
aDC
cMono
DC1-like aDC
DC1
iaDC
CXCL9+ cMono
Macrophage
ncMono
Stress DC2
Stress cMono
preDC
pDC
DC2
Cycling
C1QA+ DC2
AIRE+ aDC
Pearson's Correlation of Cluster Abundance
G
SLAMF7
CD274
CD86
ITGAX
TNFSF10
CD1C
aDC
DC1-Like aDC
iaDC
AIRE+ aDC
Fraction (%)
Normalised Expression
H
LN018 CCR7+ DC:
iaDC 3.23
94.5 aDC
LN043 CCR7+ DC:
iaDC 48.0
44.1 aDC
CD319 - PE/Dazzle-594
CD274 - PE/Fire-810
I
CD319 (SLAMF7)
CD274
CD86
CD11c (ITGAX)
TRAIL (TNFSF10)
CD1c
gMFI (x1000)
aDC iaDC

**Figure S3**

**A**

| | Acutely Inflamed | Chronically Inflamed | Control | Lymphoma | Reactive | Metastatic | Tonsillitis |
|---|---|---|---|---|---|---|---|
| This Study | 2 | 4 | | 1 | | | |
| Conde *et al.*, (2022) | | | 11 | | | | |
| Aoki *et al.*, (2020) | | | | 22 | 5 | | |
| Stewart *et al.*, (2023) | | | 7 | 2 | | | |
| Roider *et al.*, (2023) | | | | 9 | 3 | | |
| Steen *et al.*, (2021) | | | 1 | 6 | | | |
| Han *et al.*, (2022) | | | | 20 | 3 | | |
| Kim *et al.*, (2020) | | | 10 | | | 7 | |
| Pal *et al.*, (2021) | | | | | | 4 | |
| Quah *et al.*, (2023) | | | | | | 7 | |
| Choi *et al.*, (2023) | | | | | | 4 | |
| Xu *et al.*, (2021) | | | | | | 10 | |
| Massoni *et al.*, (2024) | | | | | | | 11 |

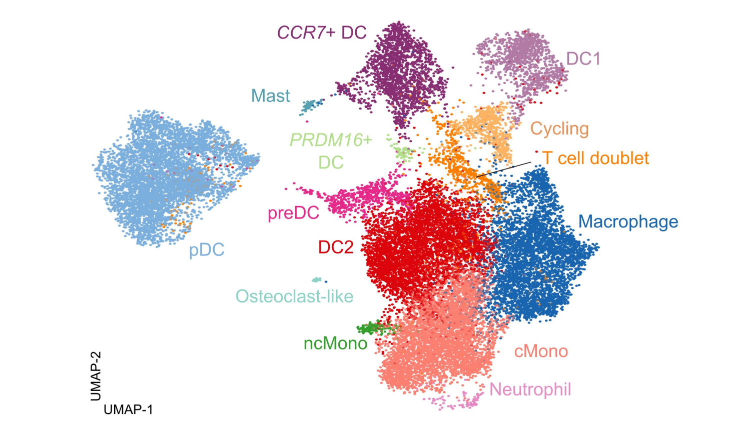


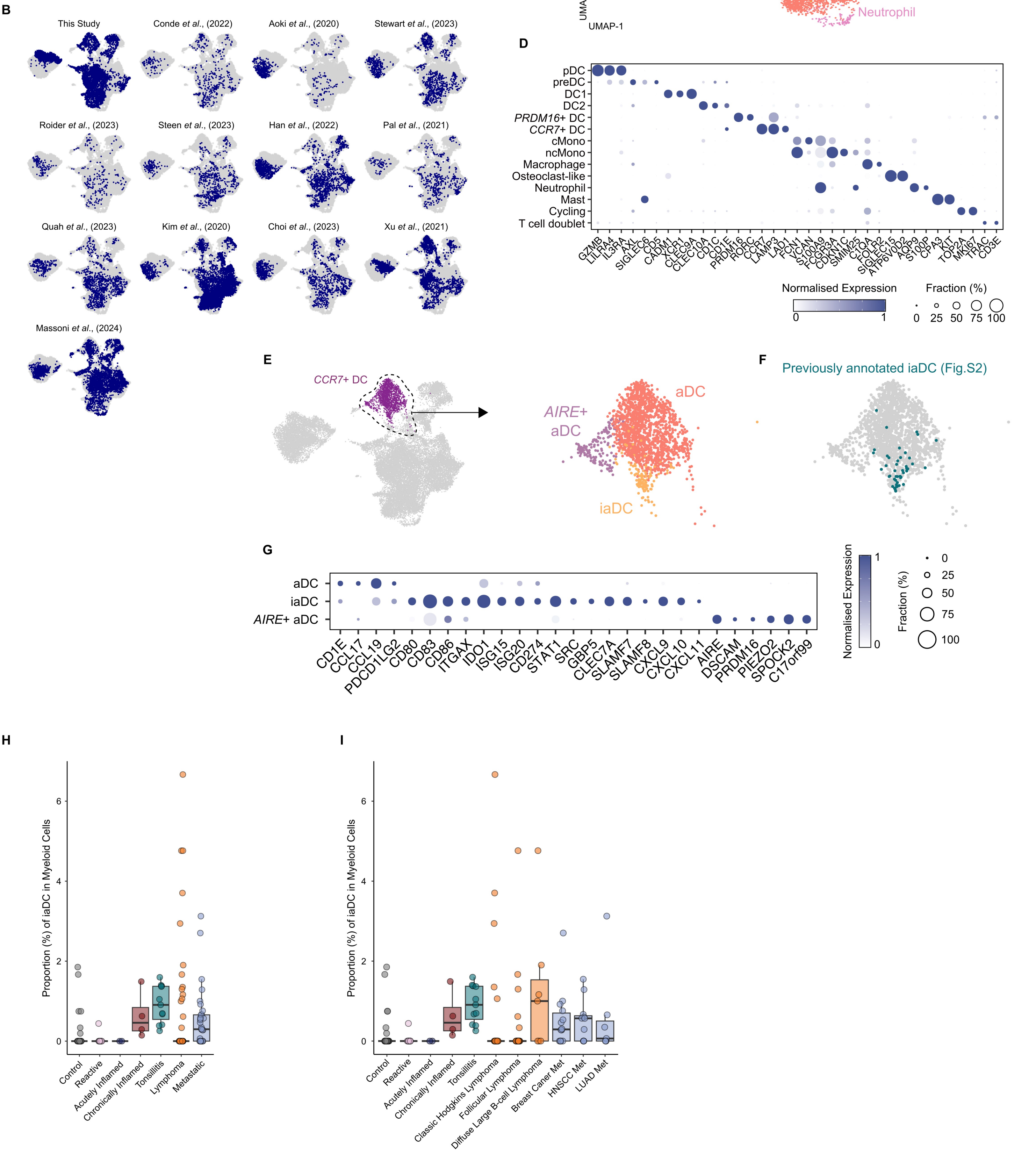

#### Figure S4

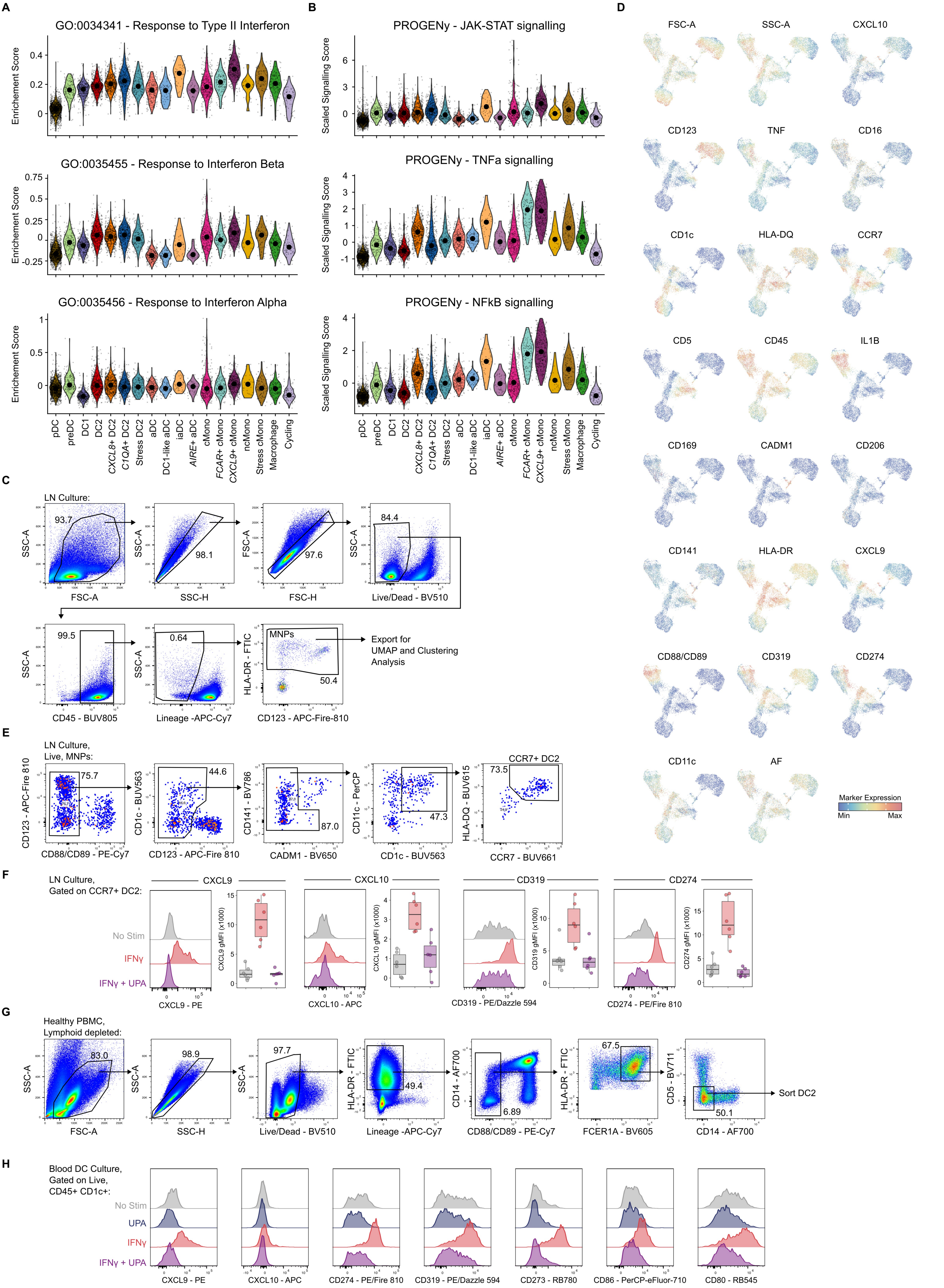
A
GO:0034341 - Response to Type II Interferon
GO:0035455 - Response to Interferon Beta
GO:0035456 - Response to Interferon Alpha
Enrichement Score
pDC
preDC
DC1
DC2
CXCL8+ DC2
C1QA+ DC2
Stress DC2
aDC
DC1-like aDC
iaDC
AIRE+ aDC
cMono
FCAR+ cMono
CXCL9+ cMono
ncMono
Stress cMono
Macrophage
Cycling
B
PROGENy - JAK-STAT signalling
PROGENy - TNFa signalling
PROGENy - NFkB signalling
Scaled Signalling Score
D
FSC-A
SSC-A
CXCL10
CD123
TNF
CD16
CD1c
HLA-DQ
CCR7
CD5
CD45
IL1B
CD169
CADM1
CD206
CD141
HLA-DR
CXCL9
CD88/CD89
CD319
CD274
CD11c
AF
Marker Expression
Min
Max
C
LN Culture:
FSC-A
SSC-H
FSC-H
Live/Dead - BV510
CD45 - BUV805
Lineage -APC-Cy7
CD123 - APC-Fire-810
HLA-DR - FTIC
MNPs
Export for
UMAP and Clustering
Analysis
E
LN Culture,
Live, MNPs:
CD123 - APC-Fire 810
CD88/CD89 - PE-Cy7
CD1c - BUV563
CD141 - BV786
CADM1 - BV650
CD11c - PerCP
CD1c - BUV563
HLA-DQ - BUV615
CCR7 - BUV661
CCR7+ DC2
F
LN Culture,
Gated on CCR7+ DC2:
CXCL9
CXCL10
CD319
CD274
No Stim
IFNγ
IFNγ + UPA
CXCL9 gMFI (x1000)
CXCL10 gMFI (x1000)
CD319 gMFI (x1000)
CD274 gMFI (x1000)
CXCL9 - PE
CXCL10 - APC
CD319 - PE/Dazzle 594
CD274 - PE/Fire 810
G
Healthy PBMC,
Lymphoid depleted:
FSC-A
SSC-H
Live/Dead - BV510
Lineage -APC-Cy7
CD14 - AF700
CD88/CD89 - PE-Cy7
FCER1A - BV605
CD5 - BV711
CD14 - AF700
Sort DC2
H
Blood DC Culture,
Gated on Live,
CD45+ CD1c+:
No Stim
UPA
IFNγ
IFNγ + UPA
CXCL9 - PE
CXCL10 - APC
CD274 - PE/Fire 810
CD319 - PE/Dazzle 594
CD273 - RB780
CD86 - PerCP-eFluor-710
CD80 - RB545

**Figure S5**

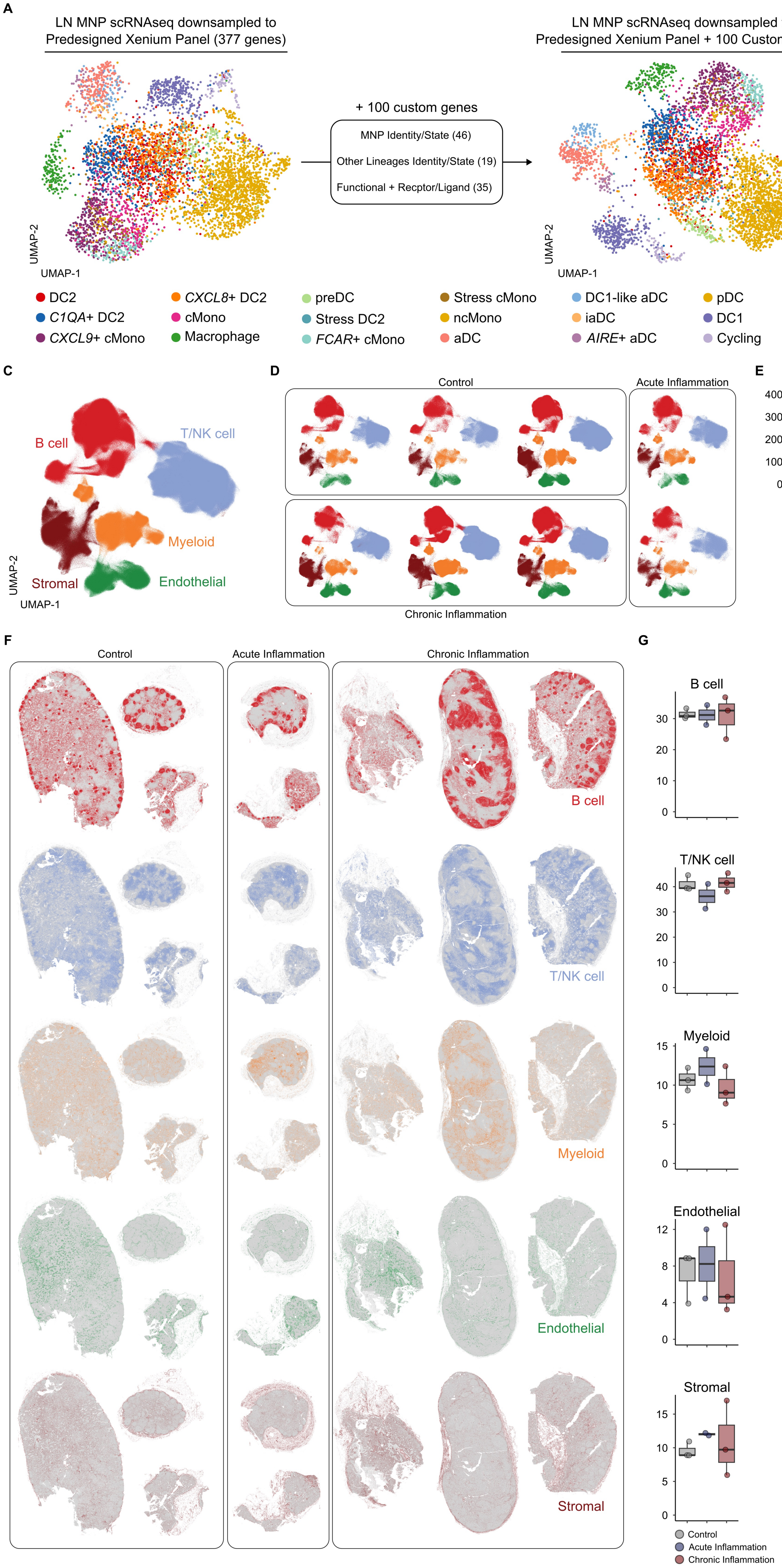
A
LN MNP scRNAseq downsampled to Predesigned Xenium Panel (377 genes)
+ 100 custom genes
MNP Identity/State (46)
Other Lineages Identity/State (19)
Functional + Recptor/Ligand (35)
LN MNP scRNAseq downsampled to Predesigned Xenium Panel + 100 Custom genes
UMAP-1
UMAP-2
DC2
C1QA+ DC2
CXCL9+ cMono
CXCL8+ DC2
cMono
Macrophage
preDC
Stress DC2
FCAR+ cMono
Stress cMono
ncMono
aDC
DC1-like aDC
iaDC
AIRE+ aDC
pDC
DC1
Cycling
B
Prediction Accuracy (%)
Predesigned
Predesigned + Custom
C
B cell
T/NK cell
Myeloid
Stromal
Endothelial
D
Control
Acute Inflammation
Chronic Inflammation
E
### Genes per segmented cell
Control
Acute Inflammation
Chronic Inflammation
F
Control
Acute Inflammation
Chronic Inflammation
B cell
T/NK cell
Myeloid
Endothelial
Stromal
G
B cell
T/NK cell
Myeloid
Endothelial
Stromal
Control
Acute Inflammation
Chronic Inflammation

**Figure S6**

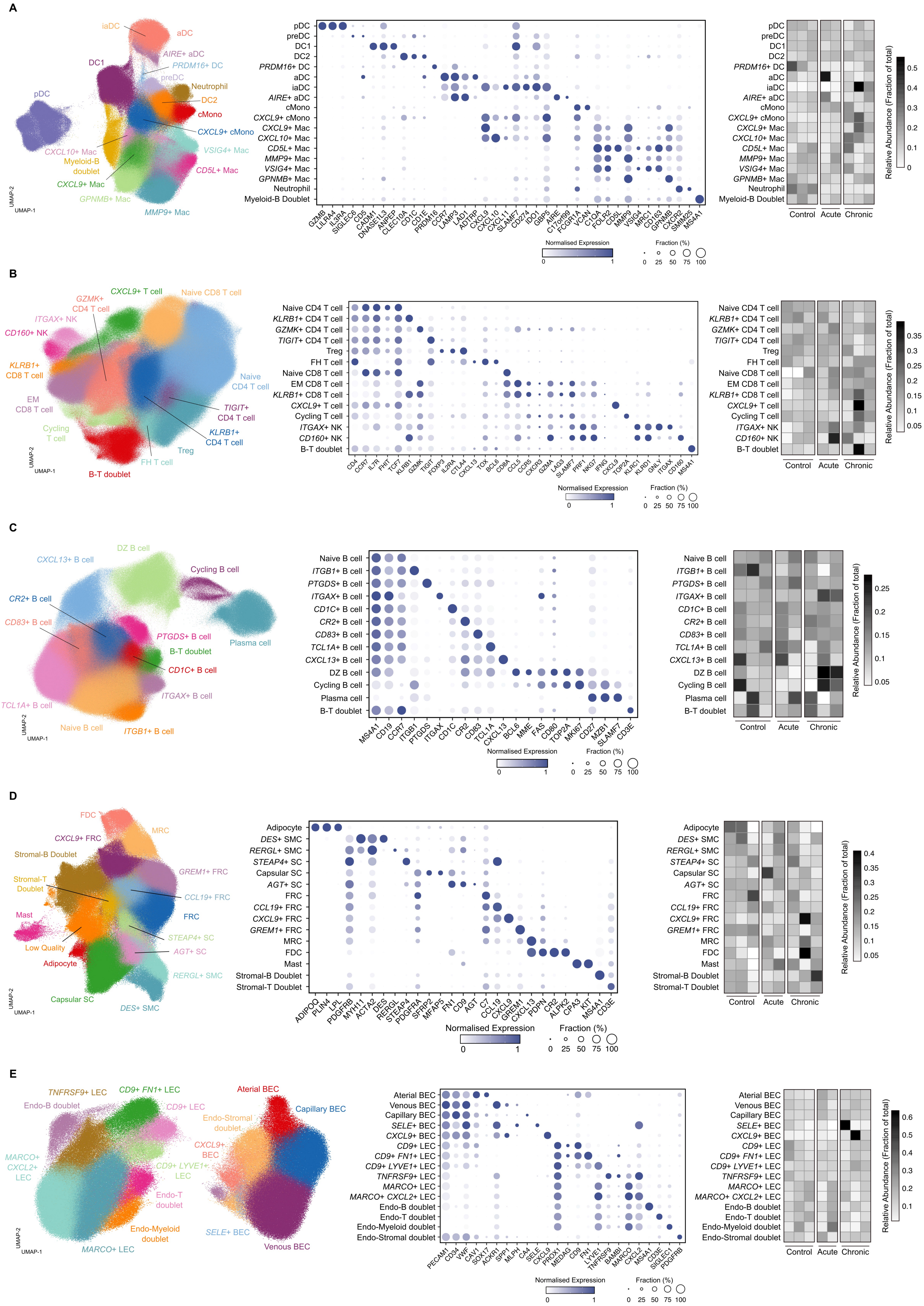

A
iaDC
aDC
AIRE+ aDC
PRDM16+ DC
preDC
DC1
pDC
Neutrophil
DC2
cMono
CXCL9+ cMono
CXCL10+ Mac
Myeloid-B doublet
CXCL9+ Mac
VSIG4+ Mac
CD5L+ Mac
GPNMB+ Mac
MMP9+ Mac
UMAP-2
UMAP-1
pDC
preDC
DC1
DC2
PRDM16+ DC
aDC
iaDC
AIRE+ aDC
cMono
CXCL9+ cMono
CXCL9+ Mac
CXCL10+ Mac
CD5L+ Mac
MMP9+ Mac
VSIG4+ Mac
GPNMB+ Mac
Neutrophil
Myeloid-B Doublet
GZMB
LILRA4
IL3RA
SIGLEC6
CD5
CADM1
DNASE1L3
ANPEP
CLEC10A
CD1C
CD1E
PRDM16
CCR7
LAMP3
LAD1
ADTRP
CXCL9
CXCL10
CXCL11
SLAMF7
CD274
IDO1
GBP5
AIRE
C17orf99
FCGR1A
VCAN
C1QA
FOLR2
CD5L
MMP9
VSIG4
MRC1
CD163
GPNMB
CXCR2
SMIM25
MS4A1
Normalised Expression
0
1
Fraction (%)
0 25 50 75 100
Control
Acute
Chronic
Relative Abundance (Fraction of total)
0.5
0.4
0.3
0.2
0.1
0
B
GZMK+ CD4 T cell
CXCL9+ T cell
Naive CD8 T cell
ITGAX+ NK
CD160+ NK
KLRB1+ CD8 T cell
EM CD8 T cell
Naive CD4 T cell
TIGIT+ CD4 T cell
Cycling T cell
KLRB1+ CD4 T cell
Treg
FH T cell
B-T doublet
UMAP-2
UMAP-1
Naive CD4 T cell
KLRB1+ CD4 T cell
GZMK+ CD4 T cell
TIGIT+ CD4 T cell
Treg
FH T cell
Naive CD8 T cell
EM CD8 T cell
KLRB1+ CD8 T cell
CXCL9+ T cell
Cycling T cell
ITGAX+ NK
CD160+ NK
B-T doublet
CD4
CCR7
IL7R
FHIT
TCF7
KLRB1
GZMK
TIGIT
FOXP3
IL2RA
CTLA4
CXCL13
TOX
BCL6
CD8A
CCL5
CCR5
CXCR3
GZMA
LAG3
SLAMF7
PRF1
NKG7
IFNG
CXCL9
TOP2A
KLRC1
KLRD1
GNLY
ITGAX
CD160
MS4A1
Normalised Expression
0
1
Fraction (%)
0 25 50 75 100
Control
Acute
Chronic
Relative Abundance (Fraction of total)
0.35
0.3
0.25
0.2
0.15
0.1
0.05
C
CXCL13+ B cell
DZ B cell
Cycling B cell
CR2+ B cell
CD83+ B cell
PTGDS+ B cell
B-T doublet
Plasma cell
CD1C+ B cell
ITGAX+ B cell
TCL1A+ B cell
Naive B cell
ITGB1+ B cell
UMAP-2
UMAP-1
Naive B cell
ITGB1+ B cell
PTGDS+ B cell
ITGAX+ B cell
CD1C+ B cell
CR2+ B cell
CD83+ B cell
TCL1A+ B cell
CXCL13+ B cell
DZ B cell
Cycling B cell
Plasma cell
B-T doublet
MS4A1
CD19
CCR7
ITGB1
PTGDS
ITGAX
CD1C
CR2
CD83
TCL1A
CXCL13
BCL6
MME
FAS
CD80
TOP2A
MKI67
CD27
MZB1
SLAMF7
CD3E
Normalised Expression
0
1
Fraction (%)
0 25 50 75 100
Control
Acute
Chronic
Relative Abundance (Fraction of total)
0.25
0.2
0.15
0.1
0.05
D
FDC
MRC
CXCL9+ FRC
Stromal-B Doublet
GREM1+ FRC
Stromal-T Doublet
CCL19+ FRC
Mast
FRC
STEAP4+ SC
Low Quality
AGT+ SC
Adipocyte
RERGL+ SMC
Capsular SC
DES+ SMC
UMAP-2
UMAP-1
Adipocyte
DES+ SMC
RERGL+ SMC
STEAP4+ SC
Capsular SC
AGT+ SC
FRC
CCL19+ FRC
CXCL9+ FRC
GREM1+ FRC
MRC
FDC
Mast
Stromal-B Doublet
Stromal-T Doublet
ADIPOQ
PLIN4
LPL
PDGFRB
MYH11
ACTA2
DES
RERGL
STEAP4
PDGFRA
SFRP2
MFAP5
FN1
CD9
AGT
C7
CCL19
CXCL9
GREM1
CXCL13
PDPN
CR2
ALPK2
CPA3
KIT
MS4A1
CD3E
Normalised Expression
0
1
Fraction (%)
0 25 50 75 100
Control
Acute
Chronic
Relative Abundance (Fraction of total)
0.4
0.35
0.3
0.25
0.2
0.15
0.1
0.05
E
TNFRSF9+ LEC
CD9+ FN1+ LEC
Aterial BEC
Endo-B doublet
CD9+ LEC
Capillary BEC
Endo-Stromal doublet
MARCO+ CXCL2+ LEC
CXCL9+ BEC
CD9+ LYVE1+ LEC
Endo-T doublet
Endo-Myeloid doublet
SELE+ BEC
MARCO+ LEC
Venous BEC
UMAP-2
UMAP-1
Aterial BEC
Venous BEC
Capillary BEC
SELE+ BEC
CXCL9+ BEC
CD9+ LEC
CD9+ FN1+ LEC
CD9+ LYVE1+ LEC
TNFRSF9+ LEC
MARCO+ LEC
MARCO+ CXCL2+ LEC
Endo-B doublet
Endo-T doublet
Endo-Myeloid doublet
Endo-Stromal doublet
PECAM1
CD34
VWF
CAV1
SOX17
ACKR1
SPP1
MLPH
CA4
SELE
CXCL9
PROX1
MEDAG
CD9
FN1
LYVE1
TNFRSF9
BAMBI
MARCO
CXCL2
MS4A1
CD3E
SIGLEC1
PDGFRB
Normalised Expression
0
1
Fraction (%)
0 25 50 75 100
Control
Acute
Chronic
Relative Abundance (Fraction of total)
0.6
0.5
0.4
0.3
0.2
0.1

**Figure S7**

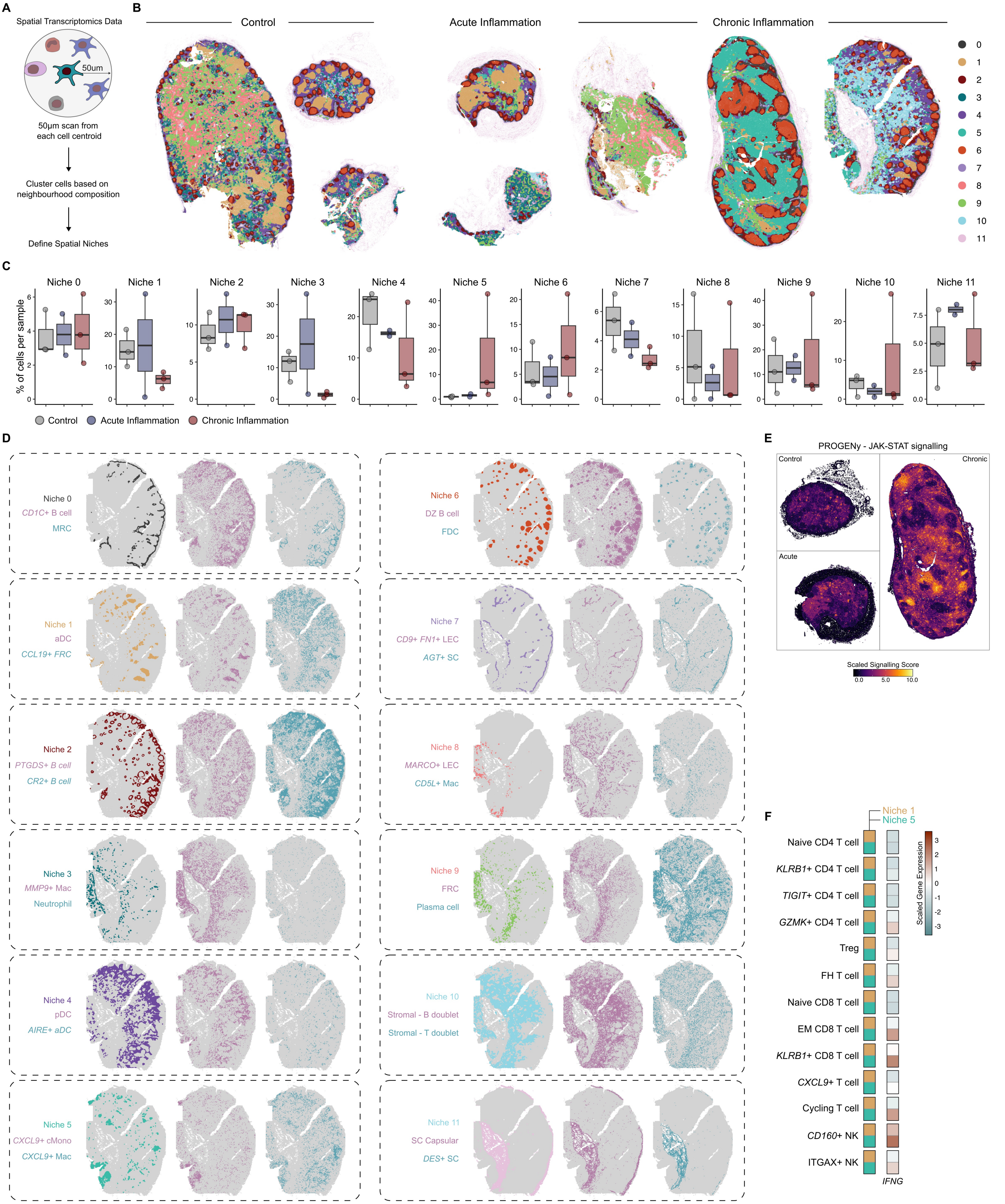

A
Spatial Transcriptomics Data
50um
50µm scan from each cell centroid
Cluster cells based on neighbourhood composition
Define Spatial Niches
B
Control
Acute Inflammation
Chronic Inflammation
0
1
2
3
4
5
6
7
8
9
10
11
C
Niche 0
Niche 1
Niche 2
Niche 3
Niche 4
Niche 5
Niche 6
Niche 7
Niche 8
Niche 9
Niche 10
Niche 11
% of cells per sample
Control
Acute Inflammation
Chronic Inflammation
D
Niche 0
CD1C+ B cell
MRC
Niche 1
aDC
CCL19+ FRC
Niche 2
PTGDS+ B cell
CR2+ B cell
Niche 3
MMP9+ Mac
Neutrophil
Niche 4
pDC
AIRE+ aDC
Niche 5
CXCL9+ cMono
CXCL9+ Mac
Niche 6
DZ B cell
FDC
Niche 7
CD9+ FN1+ LEC
AGT+ SC
Niche 8
MARCO+ LEC
CD5L+ Mac
Niche 9
FRC
Plasma cell
Niche 10
Stromal - B doublet
Stromal - T doublet
Niche 11
SC Capsular
DES+ SC
E
PROGENy - JAK-STAT signalling
Control
Chronic
Acute
Scaled Signalling Score
0.0
5.0
10.0
F
Niche 1
Niche 5
Naive CD4 T cell
KLRB1+ CD4 T cell
TIGIT+ CD4 T cell
GZMK+ CD4 T cell
Treg
FH T cell
Naive CD8 T cell
EM CD8 T cell
KLRB1+ CD8 T cell
CXCL9+ T cell
Cycling T cell
CD160+ NK
ITGAX+ NK
IFNG
Scaled Gene Expression

Figure S8

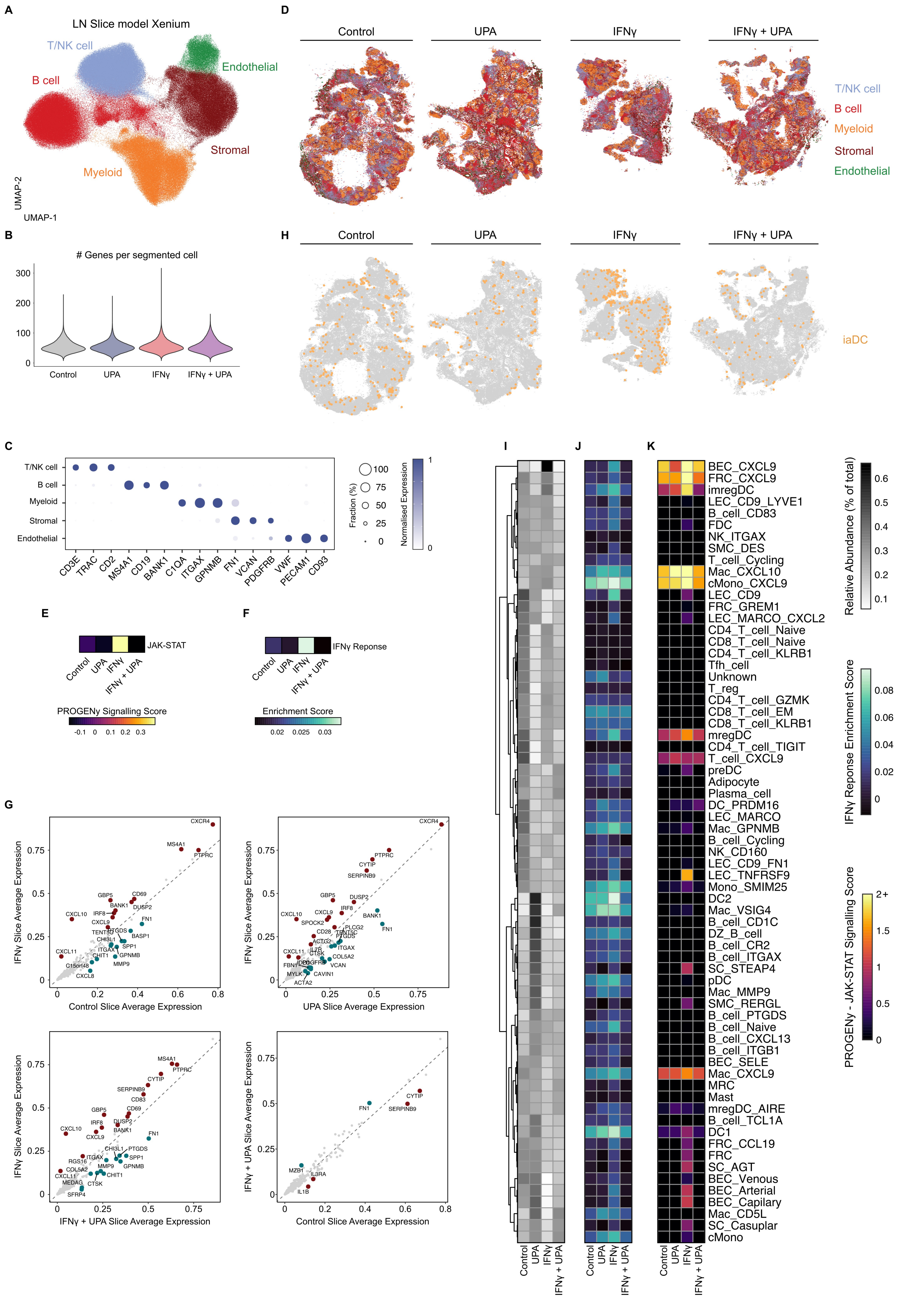

A
LN Slice model Xenium
T/NK cell
B cell
Endothelial
Stromal
Myeloid
UMAP-2
UMAP-1
B
### Genes per segmented cell
Control
UPA
IFNγ
IFNγ + UPA
C
T/NK cell
B cell
Myeloid
Stromal
Endothelial
CD3E
TRAC
CD2
MS4A1
CD19
BANK1
C1QA
ITGAX
GPNMB
FN1
VCAN
PDGFRB
VWF
PECAM1
CD93
Fraction (%)
Normalised Expression
D
Control
UPA
IFNγ
IFNγ + UPA
H
iaDC
E
JAK-STAT
PROGENy Signalling Score
F
IFNγ Reponse
Enrichment Score
G
IFNγ Slice Average Expression
Control Slice Average Expression
UPA Slice Average Expression
IFNγ + UPA Slice Average Expression
IFNγ + UPA Slice Average Expression
Control Slice Average Expression
CXCR4
MS4A1
PTPRC
CYTIP
SERPINB9
GBP5
CD69
DUSP2
BANK1
IRF8
CXCL10
CXCL9
FN1
CXCL11
SPP1
GPNMB
MMP9
FN1
MZB1
IL3RA
IL1B
I
J
K
BEC_CXCL9
FRC_CXCL9
imregDC
LEC_CD9_LYVE1
B_cell_CD83
FDC
NK_ITGAX
SMC_DES
T_cell_Cycling
Mac_CXCL10
cMono_CXCL9
LEC_CD9
FRC_GREM1
LEC_MARCO_CXCL2
CD4_T_cell_Naive
CD8_T_cell_Naive
CD4_T_cell_KLRB1
Tfh_cell
Unknown
T_reg
CD4_T_cell_GZMK
CD8_T_cell_EM
CD8_T_cell_KLRB1
mregDC
CD4_T_cell_TIGIT
T_cell_CXCL9
preDC
Adipocyte
Plasma_cell
DC_PRDM16
LEC_MARCO
Mac_GPNMB
B_cell_Cycling
NK_CD160
LEC_CD9_FN1
LEC_TNFRSF9
Mono_SMIM25
DC2
Mac_VSIG4
B_cell_CD1C
DZ_B_cell
B_cell_CR2
B_cell_ITGAX
SC_STEAP4
pDC
Mac_MMP9
SMC_RERGL
B_cell_PTGDS
B_cell_Naive
B_cell_CXCL13
B_cell_ITGB1
BEC_SELE
Mac_CXCL9
MRC
Mast
mregDC_AIRE
B_cell_TCL1A
DC1
FRC_CCL19
FRC
SC_AGT
BEC_Venous
BEC_Arterial
BEC_Capilary
Mac_CD5L
SC_Casuplar
cMono
Relative Abundance (% of total)
IFNγ Reponse Enrichment Score
PROGENy - JAK-STAT Signalling Score

**Figure S9**

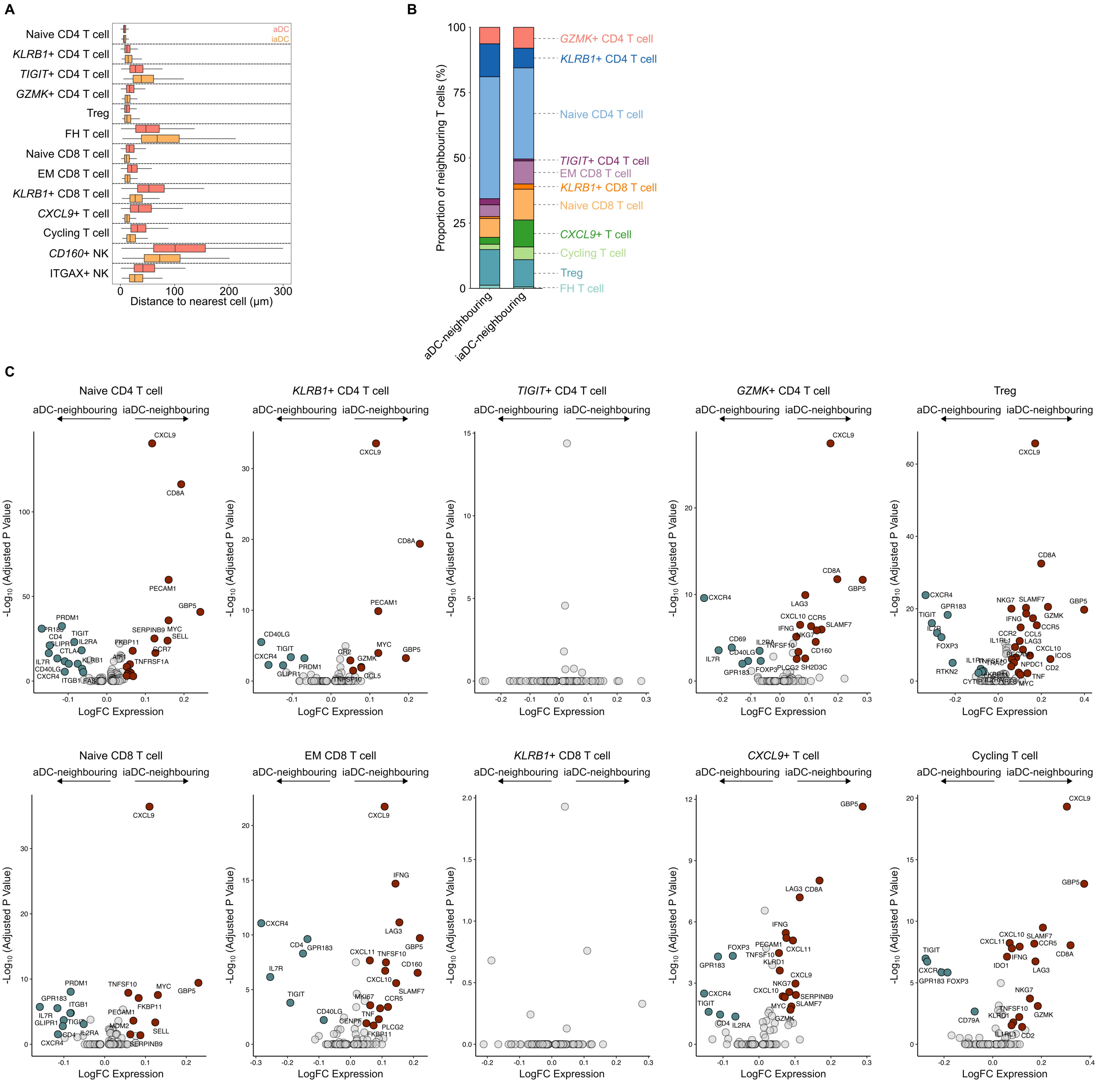
A
Naive CD4 T cell
KLRB1+ CD4 T cell
TIGIT+ CD4 T cell
GZMK+ CD4 T cell
Treg
FH T cell
Naive CD8 T cell
EM CD8 T cell
KLRB1+ CD8 T cell
CXCL9+ T cell
Cycling T cell
CD160+ NK
ITGAX+ NK
aDC
iaDC
Distance to nearest cell (μm)
B
Proportion of neighbouring T cells (%)
aDC-neighbouring
iaDC-neighbouring
GZMK+ CD4 T cell
KLRB1+ CD4 T cell
Naive CD4 T cell
TIGIT+ CD4 T cell
EM CD8 T cell
KLRB1+ CD8 T cell
Naive CD8 T cell
CXCL9+ T cell
Cycling T cell
Treg
FH T cell
C
Naive CD4 T cell
KLRB1+ CD4 T cell
TIGIT+ CD4 T cell
GZMK+ CD4 T cell
Treg
Naive CD8 T cell
EM CD8 T cell
KLRB1+ CD8 T cell
CXCL9+ T cell
Cycling T cell
aDC-neighbouring
iaDC-neighbouring
-Log10 (Adjusted P Value)
LogFC Expression

**Figure S10**

**A**

Small and Large Intestine IBD - Thomas *et al.*, 2024

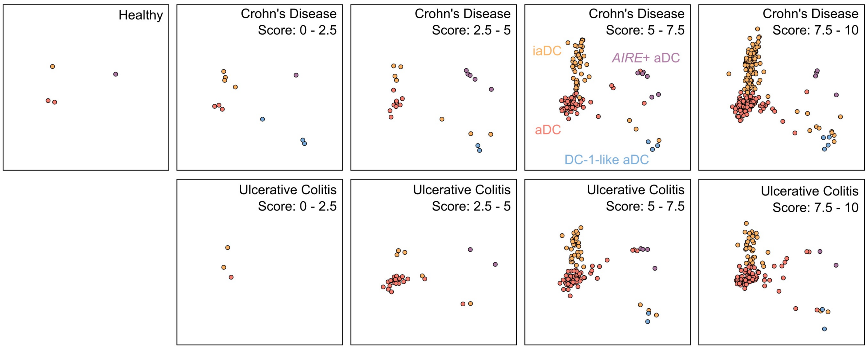


**B**

Synovial Membrane Osteoarthritis and Rheumatoid Arthritis - Zhang *et al.*, 2023

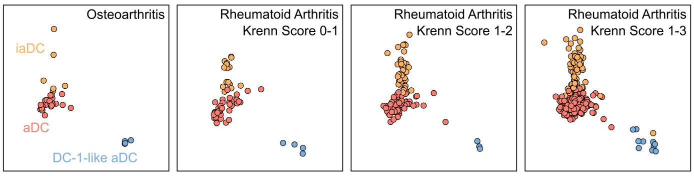


**C**

Skin Sarcoidosis - Krausgruber *et al.*, 2023

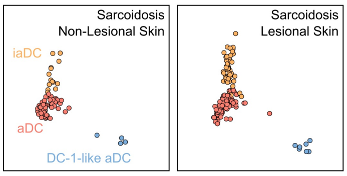

**Figure S11**

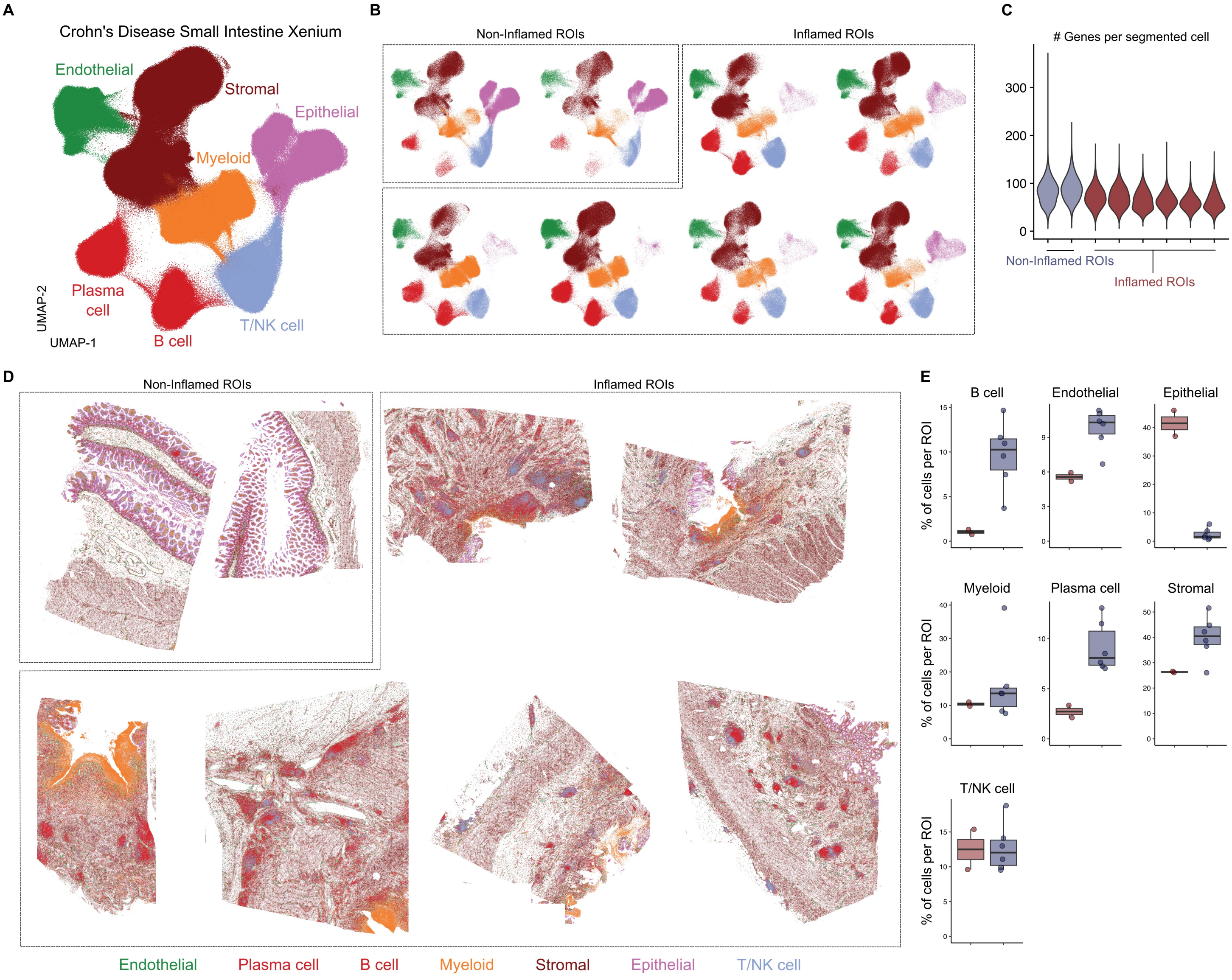

A
Crohn's Disease Small Intestine Xenium
Endothelial
Stromal
Epithelial
Myeloid
Plasma cell
B cell
T/NK cell
UMAP-2
UMAP-1
B
Non-Inflamed ROIs
Inflamed ROIs
C
### Genes per segmented cell
300
200
100
0
Non-Inflamed ROIs
Inflamed ROIs
D
Non-Inflamed ROIs
Inflamed ROIs
E
B cell
Endothelial
Epithelial
Myeloid
Plasma cell
Stromal
T/NK cell
% of cells per ROI
Endothelial
Plasma cell
B cell
Myeloid
Stromal
Epithelial
T/NK cell

**Figure S12**

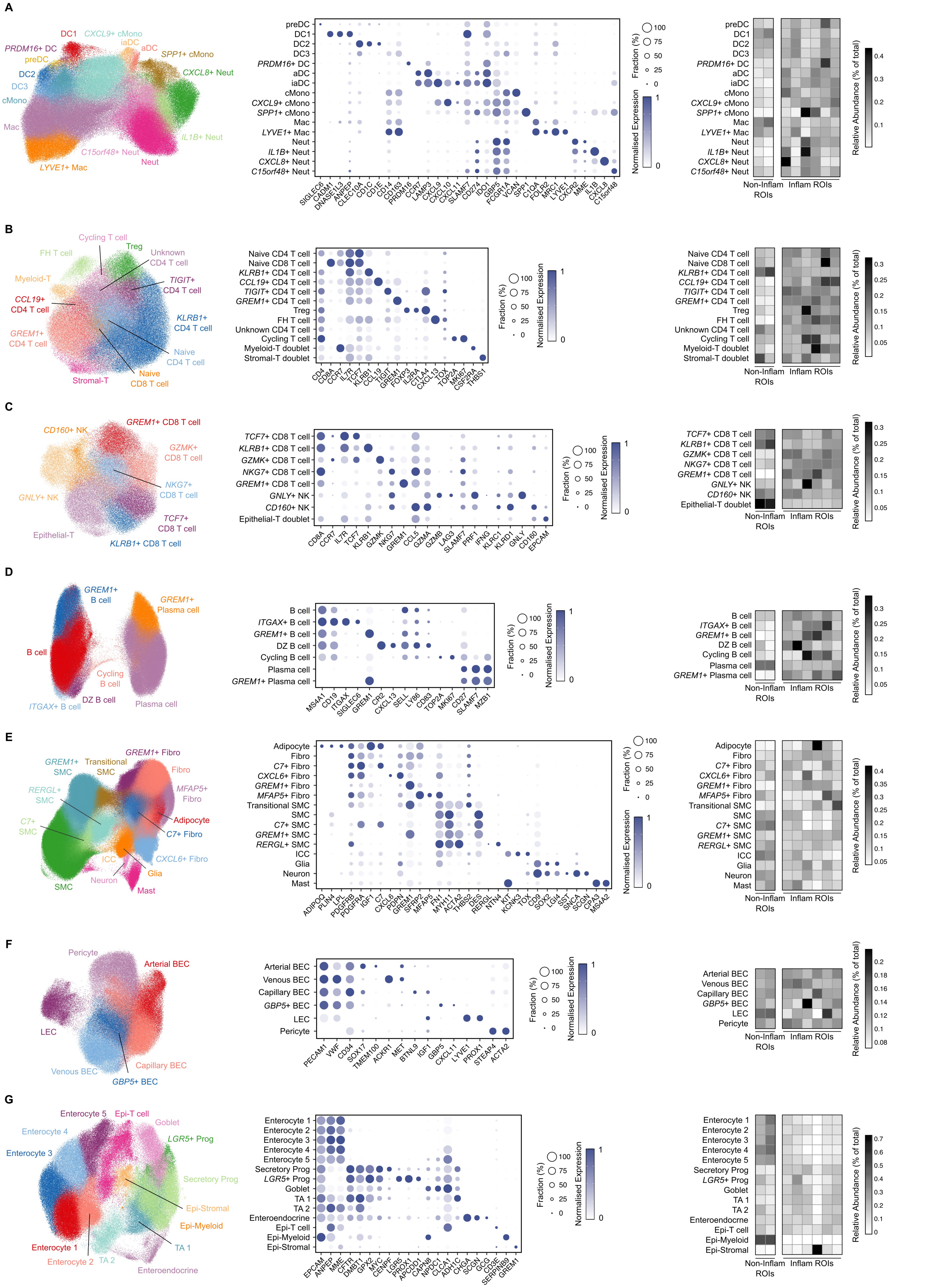
A
DC1
CXCL9+ cMono
PRDM16+ DC
iaDC
aDC
preDC
SPP1+ cMono
CXCL8+ Neut
DC2
DC3
cMono
Mac
IL1B+ Neut
C15orf48+ Neut
Neut
LYVE1+ Mac
Fraction (%)
Normalised Expression
Relative Abundance (% of total)
Non-Inflam ROIs
Inflam ROIs
B
Cycling T cell
FH T cell
Treg
Unknown CD4 T cell
Myeloid-T
TIGIT+ CD4 T cell
CCL19+ CD4 T cell
KLRB1+ CD4 T cell
GREM1+ CD4 T cell
Naive CD4 T cell
Naive CD8 T cell
Stromal-T
Myeloid-T doublet
Stromal-T doublet
C
CD160+ NK
GREM1+ CD8 T cell
GZMK+ CD8 T cell
NKG7+ CD8 T cell
GNLY+ NK
TCF7+ CD8 T cell
Epithelial-T
KLRB1+ CD8 T cell
Epithelial-T doublet
D
GREM1+ B cell
GREM1+ Plasma cell
B cell
Cycling B cell
DZ B cell
ITGAX+ B cell
Plasma cell
E
GREM1+ Fibro
GREM1+ SMC
Transitional SMC
Fibro
RERGL+ SMC
MFAP5+ Fibro
C7+ SMC
Adipocyte
C7+ Fibro
ICC
CXCL6+ Fibro
Glia
Neuron
SMC
Mast
F
Pericyte
Arterial BEC
LEC
Venous BEC
Capillary BEC
GBP5+ BEC
G
Enterocyte 1
Enterocyte 2
Enterocyte 3
Enterocyte 4
Enterocyte 5
Epi-T cell
Goblet
LGR5+ Prog
Secretory Prog
Epi-Stromal
Epi-Myeloid
TA 1
TA 2
Enteroendocrine

**Figure S13**

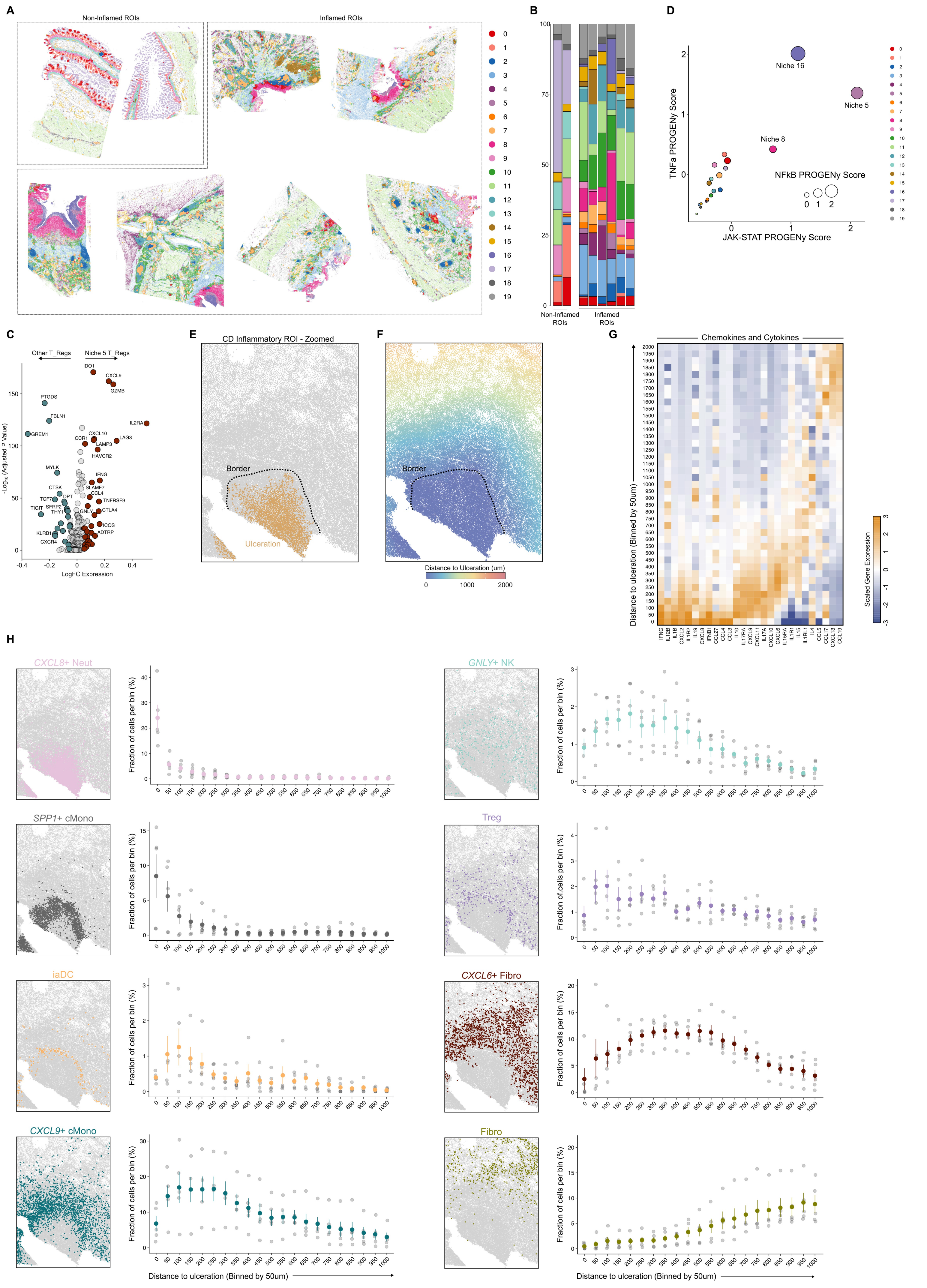
A
Non-Inflamed ROIs
Inflamed ROIs
B
Non-Inflamed ROIs
Inflamed ROIs
D
TNFa PROGENy Score
JAK-STAT PROGENy Score
Niche 16
Niche 5
Niche 8
NFkB PROGENy Score
C
Other T_Regs
Niche 5 T_Regs
-Log10 (Adjusted P Value)
LogFC Expression
IDO1
CXCL9
GZMB
PTGDS
FBLN1
IL2RA
GREM1
CXCL10
CCR1
LAG3
LAMP3
HAVCR2
MYLK
IFNG
CTSK
SLAMF7
DPT
CCL4
TNFRSF9
TCF7
TIGIT
SFRP2
THY1
GNLY
CTLA4
ICOS
KLRB1
ADTRP
CXCR4
E
CD Inflammatory ROI - Zoomed
Border
Ulceration
F
Border
Distance to Ulceration (um)
G
Chemokines and Cytokines
Distance to ulceration (Binned by 50um)
Scaled Gene Expression
IFNG
IL12B
IL1B
CXCL2
IL1R2
IL19
CXCL8
IFNB1
CCL27
CCL4
CCL3
IL10
IL17RA
CXCL9
CXCL11
IL17A
CXCL10
CXCL6
IL15RA
IL1R1
IL15
IL1RL1
IL4
CCL5
CCL17
CXCL13
CCL19
H
CXCL8+ Neut
GNLY+ NK
SPP1+ cMono
Treg
iaDC
CXCL6+ Fibro
CXCL9+ cMono
Fibro
Fraction of cells per bin (%)
Distance to ulceration (Binned by 50um)

**Figure S14**

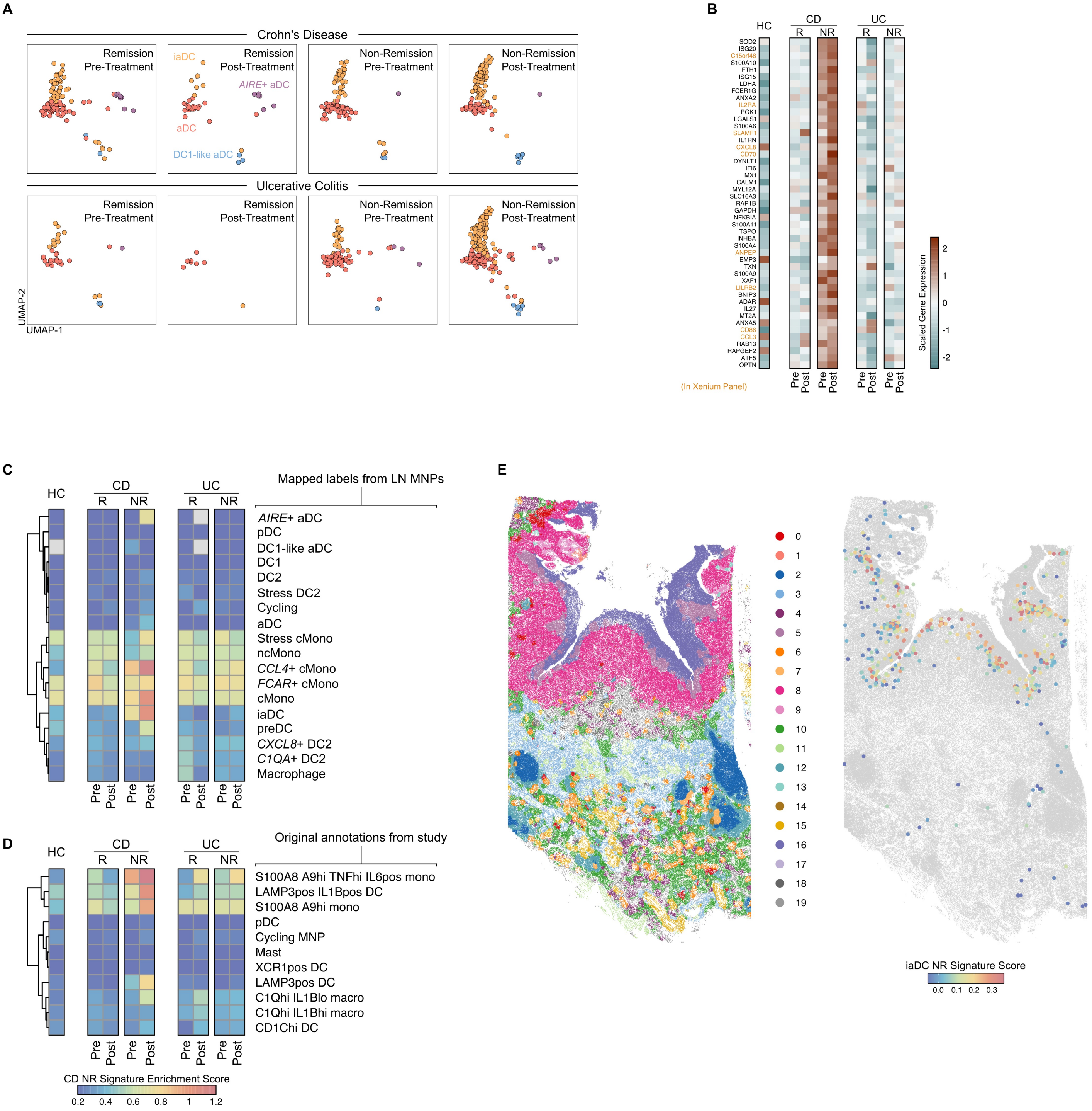
A
Crohn's Disease
Remission Pre-Treatment
Remission Post-Treatment
Non-Remission Pre-Treatment
Non-Remission Post-Treatment
iaDC
AIRE+ aDC
aDC
DC1-like aDC
Ulcerative Colitis
UMAP-2
UMAP-1
B
HC
CD
UC
R
NR
SOD2
ISG20
C15orf48
S100A10
FTH1
ISG15
LDHA
FCER1G
ANXA2
IL2RA
PGK1
LGALS1
S100A6
SLAMF1
IL1RN
CXCL8
CD70
DYNLT1
IFI6
MX1
CALM1
MYL12A
SLC16A3
RAP1B
GAPDH
NFKBIA
S100A11
TSPO
INHBA
S100A4
ANPEP
EMP3
TXN
S100A9
XAF1
LILRB2
BNIP3
ADAR
IL27
MT2A
ANXA5
CD86
CCL3
RAB13
RAPGEF2
ATF5
OPTN
(In Xenium Panel)
Pre
Post
Scaled Gene Expression
C
Mapped labels from LN MNPs
AIRE+ aDC
pDC
DC1-like aDC
DC1
DC2
Stress DC2
Cycling
aDC
Stress cMono
ncMono
CCL4+ cMono
FCAR+ cMono
cMono
iaDC
preDC
CXCL8+ DC2
C1QA+ DC2
Macrophage
D
Original annotations from study
S100A8 A9hi TNFhi IL6pos mono
LAMP3pos IL1Bpos DC
S100A8 A9hi mono
pDC
Cycling MNP
Mast
XCR1pos DC
LAMP3pos DC
C1Qhi IL1Blo macro
C1Qhi IL1Bhi macro
CD1Chi DC
CD NR Signature Enrichment Score
0.2 0.4 0.6 0.8 1 1.2
E
0 1 2 3 4 5 6 7 8 9 10 11 12 13 14 15 16 17 18 19
iaDC NR Signature Score
0.0 0.1 0.2 0.3

**Figure S15**

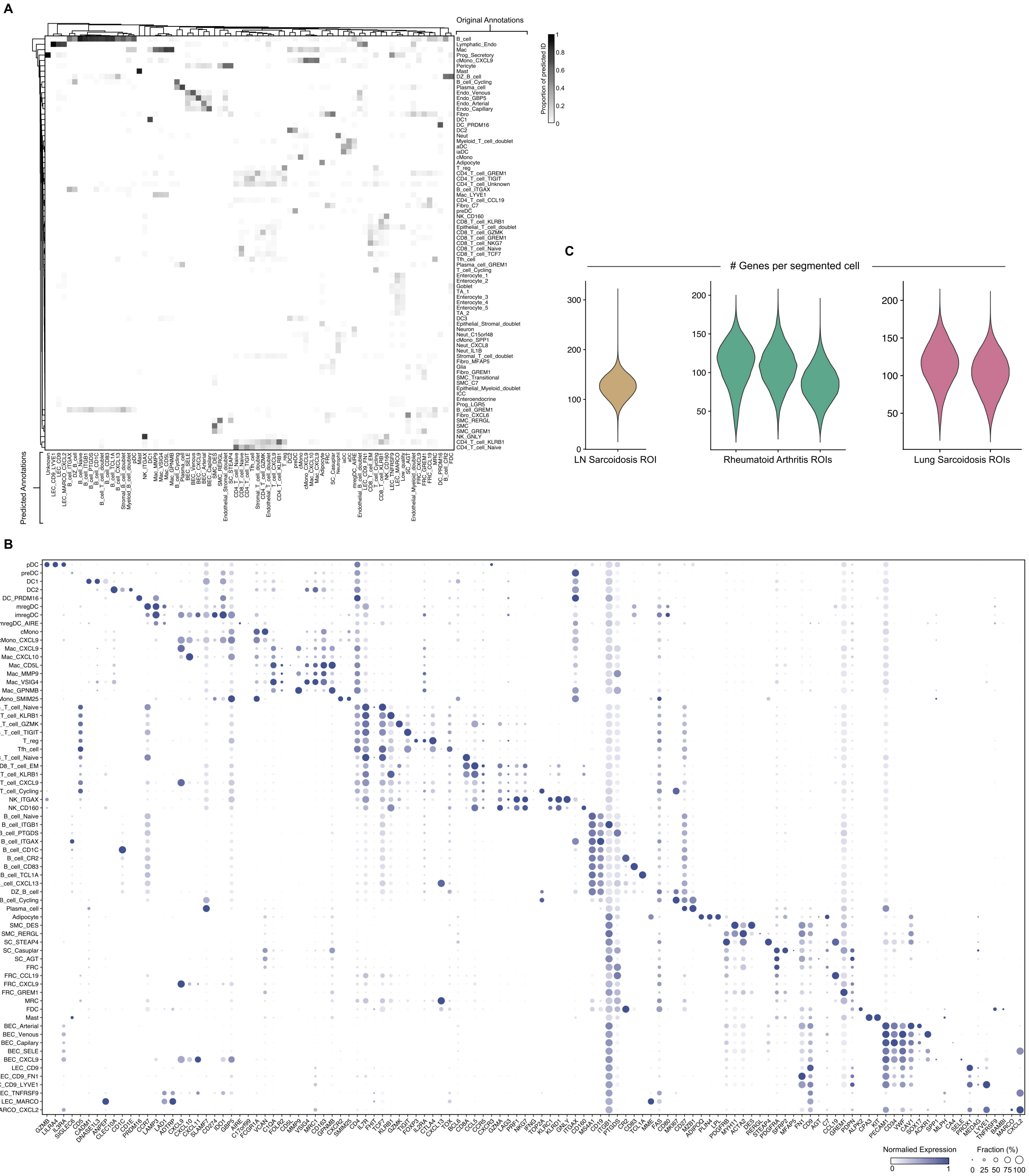
A
Original Annotations
Predicted Annotations
Proportion of predicted ID
1
0.8
0.6
0.4
0.2
0
B_cell
Lymphatic_Endo
Mac
Prog_Secretory
cMono_CXCL9
Pericyte
Mast
DZ_B_cell
B_cell_Cycling
Plasma_cell
Endo_Venous
Endo_GBP5
Endo_Arterial
Endo_Capillary
Fibro
DC1
DC_PRDM16
DC2
Neut
Myeloid_T_cell_doublet
aDC
iaDC
cMono
Adipocyte
T_reg
CD4_T_cell_GREM1
CD4_T_cell_TIGIT
CD4_T_cell_Unknown
B_cell_ITGAX
Mac_LYVE1
CD4_T_cell_CCL19
Fibro_C7
preDC
NK_CD160
CD8_T_cell_KLRB1
Epithelial_T_cell_doublet
CD8_T_cell_GZMK
CD8_T_cell_GREM1
CD8_T_cell_NKG7
CD8_T_cell_Naive
CD8_T_cell_TCF7
Tfh_cell
Plasma_cell_GREM1
T_cell_Cycling
Enterocyte_1
Enterocyte_2
Goblet
TA_1
Enterocyte_3
Enterocyte_4
Enterocyte_5
TA_2
DC3
Epithelial_Stromal_doublet
Neuron
Neut_C15orf48
cMono_SPP1
Neut_CXCL8
Neut_IL1B
Stromal_T_cell_doublet
Fibro_MFAP5
Glia
Fibro_GREM1
SMC_Transitional
SMC_C7
Epithelial_Myeloid_doublet
ICC
Enteroendocrine
Prog_LGR5
B_cell_GREM1
Fibro_CXCL6
SMC_RERGL
SMC
SMC_GREM1
NK_GNLY
CD4_T_cell_KLRB1
CD4_T_cell_Naive
C
### Genes per segmented cell
LN Sarcoidosis ROI
Rheumatoid Arthritis ROIs
Lung Sarcoidosis ROIs
B
pDC
preDC
DC1
DC2
DC_PRDM16
mregDC
imregDC
mregDC_AIRE
cMono
cMono_CXCL9
Mac_CXCL9
Mac_CXCL10
Mac_CD5L
Mac_MMP9
Mac_VSIG4
Mac_GPNMB
Mono_SMIM25
CD4_T_cell_Naive
CD4_T_cell_KLRB1
CD4_T_cell_GZMK
CD4_T_cell_TIGIT
T_reg
Tfh_cell
CD8_T_cell_Naive
CD8_T_cell_EM
CD8_T_cell_KLRB1
T_cell_CXCL9
T_cell_Cycling
NK_ITGAX
NK_CD160
B_cell_Naive
B_cell_ITGB1
B_cell_PTGDS
B_cell_ITGAX
B_cell_CD1C
B_cell_CR2
B_cell_CD83
B_cell_TCL1A
B_cell_CXCL13
DZ_B_cell
B_cell_Cycling
Plasma_cell
Adipocyte
SMC_DES
SMC_RERGL
SC_STEAP4
SC_Casuplar
SC_AGT
FRC
FRC_CCL19
FRC_CXCL9
FRC_GREM1
MRC
FDC
Mast
BEC_Arterial
BEC_Venous
BEC_Capilary
BEC_SELE
BEC_CXCL9
LEC_CD9
LEC_CD9_FN1
LEC_CD9_LYVE1
LEC_TNFRSF9
LEC_MARCO
LEC_MARCO_CXCL2
Normalied Expression
0
1
Fraction (%)
0 25 50 75 100
